# Pulsed transcranial photobiomodulation evokes sustained, parameter-dependent changes in EEG oscillations that run opposite to frequency entrainment

**DOI:** 10.1101/2025.05.26.656199

**Authors:** Alicia A. Mathew, Hannah Van Lankveld, Xiaole Z. Zhong, Joanna X. Chen, Reza Zomorrodi, J. Jean Chen

## Abstract

**Objectives:** Transcranial photobiomodulation (tPBM) modulates cortical activity, but how specific stimulation parameters shape the electrophysiological response remains unclear. We tested whether pulsed forehead tPBM produces frequency band-specific EEG changes, whether pulsation frequency entrains endogenous oscillations, and whether wavelength, pulsation frequency, irradiance, sex, and skin tone significantly modulate the response.

**Materials and Methods:** Forty-six healthy young adults (24M/22F, 20-32 years) each completed four 12-minute EEG recordings (256-channel; PRE-DURING-POST, 4 min each) with pulsed NIR light delivered to the right forehead. The parameter space included two wavelengths (808 nm, 1064 nm), two pulsation frequencies (10 Hz, 40 Hz), and three irradiances (100, 150, 200 mW/cm^2^). Forehead skin tone (ITA) and sex were included as biological moderators. EEG band power (delta through gamma) was expressed as percent change from a pre-stimulus baseline. Spatiotemporal cluster-based permutation tests (10,000 permutations, FWER alpha=0.05) identified significant electrode clusters. Linear mixed-effects models with backward elimination and FDR correction (q=0.05) quantified parameter and biological contributions.

**Results:** tPBM produced bilateral, bidirectional EEG changes: frontal gamma enhancement and posterior theta, alpha, and beta suppression, emerging gradually during stimulation and persisting post-stimulation. Wavelength qualitatively shaped the response: 808 nm drove broadband posterior suppression while 1064 nm selectively enhanced frontal gamma. Contrary to entrainment predictions, 10 Hz pulsation produced significantly larger gamma increases and theta reductions than 40 Hz. Sex was the most consistent biological modulator: females showed greater alpha suppression during stimulation and males showed greater posterior beta and gamma suppression. Irradiance and skin tone were not significant.

**Conclusions:** Pulsed forehead tPBM produces robust, parameter-dependent EEG changes that are not consistent with neural entrainment. Wavelength determines the character of the cortical response, pulsation frequency modulates its magnitude, potentially through photobiological rather than oscillatory mechanisms, and sex is a significant biological moderator. These findings contribute toward a human-specific foundation for informing optical tPBM parameter selection.

## 1 INTRODUCTION

Photobiomodulation (PBM), or low-level light therapy, is a non-invasive brain stimulation technique that uses low-power near-infrared (NIR) light (∼600-1100 nm) to enhance brain metabolism and function. Transcranial photobiomodulation (tPBM) delivers this light through the scalp, where it is absorbed by mitochondrial cytochrome c oxidase (CCO) in cortical neurons.^1,2^ As the terminal enzyme of the electron transport chain, CCO drives ATP synthesis,^3^ a process inhibited by nitric oxide (NO), which competes with oxygen for binding at CCO. tPBM photodissociates NO from CCO, freeing the enzyme to bind oxygen,^4^ thereby increasing cellular energy production. The released NO can also diffuse into the surrounding vasculature, where it acts as a vasodilator and raises regional cerebral blood flow, an alternate, hemodynamic pathway through which tPBM may support brain function.^5,6^ Beyond these metabolic and vascular routes, PBM modulates ion channels, including glutamate (NMDA/AMPA) and acetylcholine receptors and voltage-gated potassium and sodium channels, both indirectly via CCO-driven cascades and, for some channels, through direct photon absorption, shaping neuronal excitability.^7^

Potentially, through these mechanisms, tPBM has been associated with improved attention, memory,^8,9^ reaction time^10^, and cognitive flexibility,^11^ and is under clinical investigation for stroke, traumatic brain injury, depression, and neurodegeneration.^12^ Yet its reliability remains constrained by poorly understood parameter dependencies.^13^

Past research has used NIR spectroscopy (NIRS), functional magnetic resonance imaging (fMRI), and electroencephalography (EEG) to investigate the brain’s physiological response to tPBM (see Review).^13^ Across these methods, tPBM has been shown to elicit local and distributed changes in CCO metabolism, hemodynamics, and neural excitability that often persist after stimulation ends.^14–20^ EEG, with its high temporal resolution, is particularly suited to characterizing these effects in real time, and prior works report changes in neural oscillations; often a relative shift toward faster rhythms, favoring better cognition, though their direction and magnitude vary considerably across protocols (see Review).^13^ Critically, most EEG studies test only one or a few parameters, leaving unclear how in vivo responses scale with specific stimulation variables. The field thus lacks a clear picture of individual parameter contributions to the dose-response relationship, essential to advancing tPBM as a precision-medicine intervention.

A further, almost unexamined dimension is pulsed light. Most tPBM studies use continuous wave delivery ^13^, yet pulsing creates an opportunity for neural entrainment: the synchronization of endogenous oscillations to a periodic external input^21,22^. This signature is well characterized for rhythmic sensory and electromagnetic stimulation^21,23–26^ but not for PBM. Tang et al. compared continuous versus pulsed light across two frequencies (40 Hz, 100 Hz) and two wavelengths (660 nm, 850 nm), finding that pulsed light raised gamma power, yet frequencies did not differ.^27^ Whether pulsed tPBM produces frequency-specific power changes at commonly used wavelengths remains unresolved. Importantly, whether 40 Hz flickering truly entrains native gamma is contested even in sensory paradigms^28^, and tPBM is mechanistically distinct – light is absorbed directly by cortical tissue rather than routed through sensory pathways. Zomorrodi et al. reported that 40 Hz tPBM increased high- and decreased low-frequency EEG power, but whether different pulsation frequencies target different rhythms remains untested.^29^ We contrast 40 Hz, the canonical gamma-band target, with 10 Hz, which falls in the alpha band, the dominant rhythm of the resting cortex and an index of attentional control^30,31^. Because entrainment should be strongest when the stimulus matches an intrinsic rhythm, we hypothesize that 10 Hz and 40 Hz pulsed tPBM will alter alpha and gamma activity, respectively.

Beyond light parameters, the effective cortical dose also depends on individual biology. Because melanin strongly absorbs NIR light,^32^ darker skin tones may attenuate the energy reaching the brain,^33^ and sex differences in baseline neural activity may further moderate the response. ^34^ These factors are rarely accounted for in tPBM-EEG studies.

We address these gaps using high-density EEG to characterize how pulsed forehead tPBM influences neural oscillations in 46 healthy young adults. Our parameter space spans two wavelengths acting through partly distinct routes: 808 nm, near CCO’s absorption peak, and 1064 nm, which scatters less and penetrates deeper but is absorbed more by water, shifting action toward thermally sensitive mechanisms such as transient receptor potential (TRP) channel gating.^35–39^ We vary two pulsation frequencies (10 Hz, 40 Hz) and three irradiances (100, 150, 200 mW/cm²),^40^ and treat skin tone (ITA) and sex as candidate biological moderators. Our aims are to (i) test whether pulsed forehead tPBM reproduces oscillatory changes reported in the literature; (ii) assess whether it entrains endogenous rhythms; (iii) map the spatiotemporal modulation by stimulation parameter; and (iv) assess modulation by differences in wavelength, pulsation frequency, irradiance, sex, and skin tone.

## 2 MATERIALS AND METHODS

### 2.1 Participants and skin tone measurements

Forty-six healthy young adults (20-32 years; 24M/22F) were recruited from Baycrest’s participant database. Exclusion criteria included neurological or physiological disorders and substance use. Ethics approval was obtained from the Baycrest Research Ethics Board and all participants provided written informed consent. Each participant’s forehead skin tone was measured using a spectrophotometer (CM-600D, Konica Minolta, Tokyo, Japan). Six readings from an 8-mm area on the right forehead yielded L* and b* values in the CIELAB color space (**Fig. 1a**), used to calculate the individual typology angle (ITA)^41^ via **Eq. 1**. Higher ITA corresponds to lighter skin and lower eumelanin.^42,43^ ITA was treated continuously in analyses but stratified into three groups for recruitment balance (**Fig. 1b**). Additional details are in supplementary **Sections S1 and S2.1**.

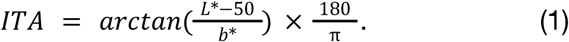

**Figure 1.**
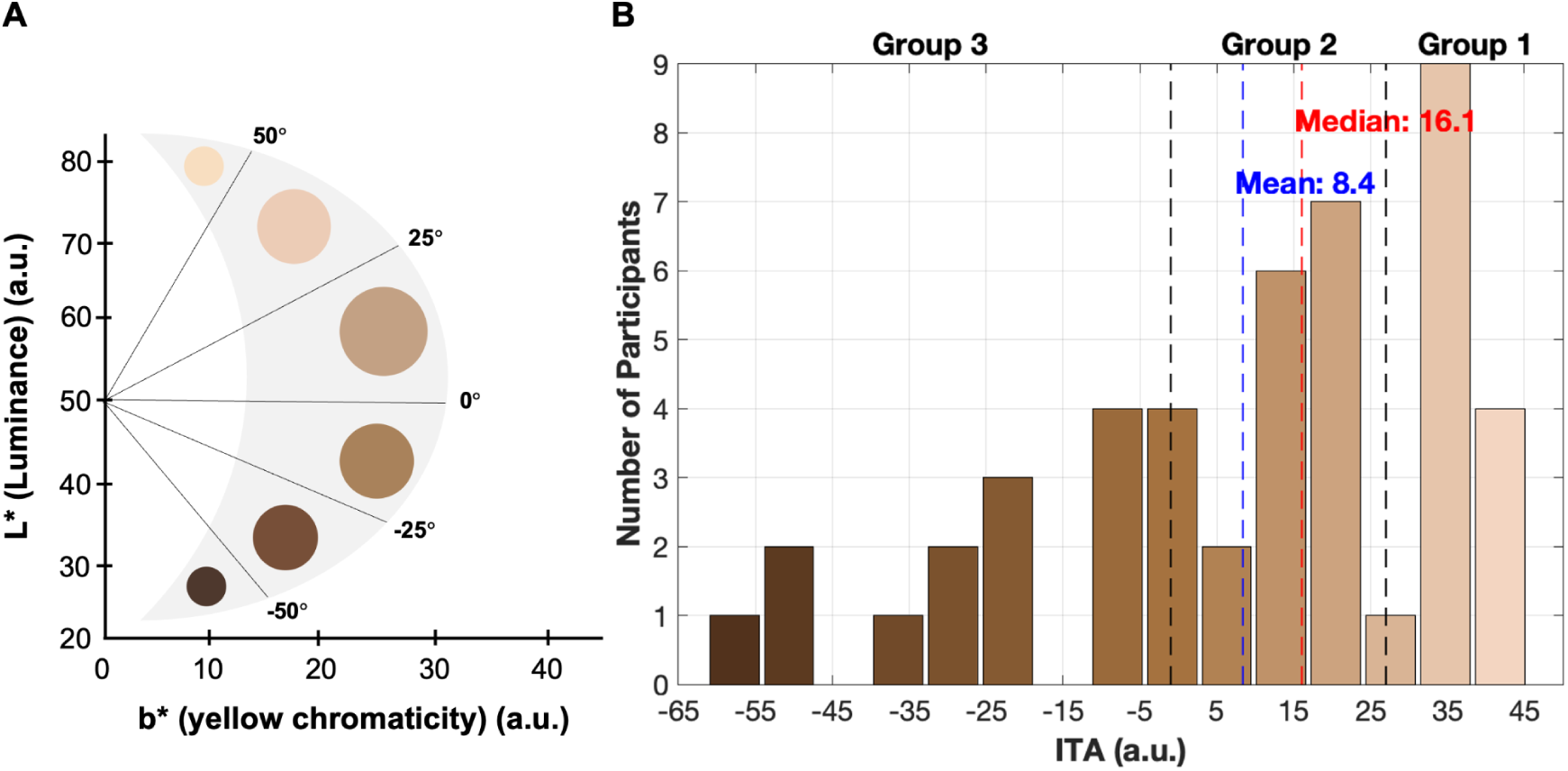
ITA as an index of skin tone. (A) ITA derived from L* and b* values in the CIELAB color space. (B) Participant distribution across ITAs (median ± range = 16.1±106.7). Mean and median values shown in blue and red, and group boundaries in black.

### 2.2 Experimental design and tPBM protocol

Participants were randomly assigned to one of three protocols determining the light parameters for each of four EEG-tPBM recordings (**Table 1**): two wavelengths (808 nm, 1064 nm), two pulsation frequencies (10 Hz, 40 Hz), and three irradiances (100, 150, 200 mW/cm^2^), chosen based on their established tissue penetration and safety profiles and their efficacy in past work.^40^

**Table 1.** Participant distribution across skin tone groups, protocols, and sexes.

| Skin Tone Group | ITA Range | Protocol 1 | Protocol 2 | Protocol 3 | Total | M / F |
| --- | --- | --- | --- | --- | --- | --- |
| Group 1 | $27^{\circ} < \text{ITA} \leq 55^{\circ}$ | 5 | 5 | 4 | 14 | 6 M / 8 F |
| Group 2 | $-1^{\circ} < \text{ITA} \leq 27^{\circ}$ | 6 | 5 | 6 | 17 | 9 M / 8 F |
| Group 3 | $\text{ITA} \leq -1^{\circ}$ | 5 | 4 | 6 | 15 | 9 M / 6 F |
| **Number of Subjects** |  | 16 | 14 | 16 | 46 | 24 M / 22 F |
| **Number of Recordings** |  | 64 | 56 | 64 | 184 |  |

Each 12-minute recording followed a PRE-DURING-POST block design (4 minutes each; **Fig. 2b**). Participants watched naturalistic-stimulus videos to minimize drowsiness and control brain state ^44^. The PRE period served as the within-subject baseline. During stimulation, NIR light was delivered to the right forehead targeting the right prefrontal cortex through a custom headpiece via a 10-meter, 400 µm fiber cable and two Class 3 laser systems (MDL-III-808-1W and MDL-III-1064-1W; courtesy of Vielight Inc., Toronto, Canada; **Fig. 2a**) at a 50% duty cycle. Light parameters were controlled from behind the participant so they remained blinded to the protocols and study design. Dosage parameters are in **Table 3**; calibration details in supplementary **Section S2**. A 22-minute break separated consecutive recordings. Finally, a separate magnetic resonance (MR) thermometry scan was collected to confirm that no measurable heating effects were produced in the targeted region, with participants also reporting no thermal sensations (supplementary **Section S3**).

**Figure 2.**
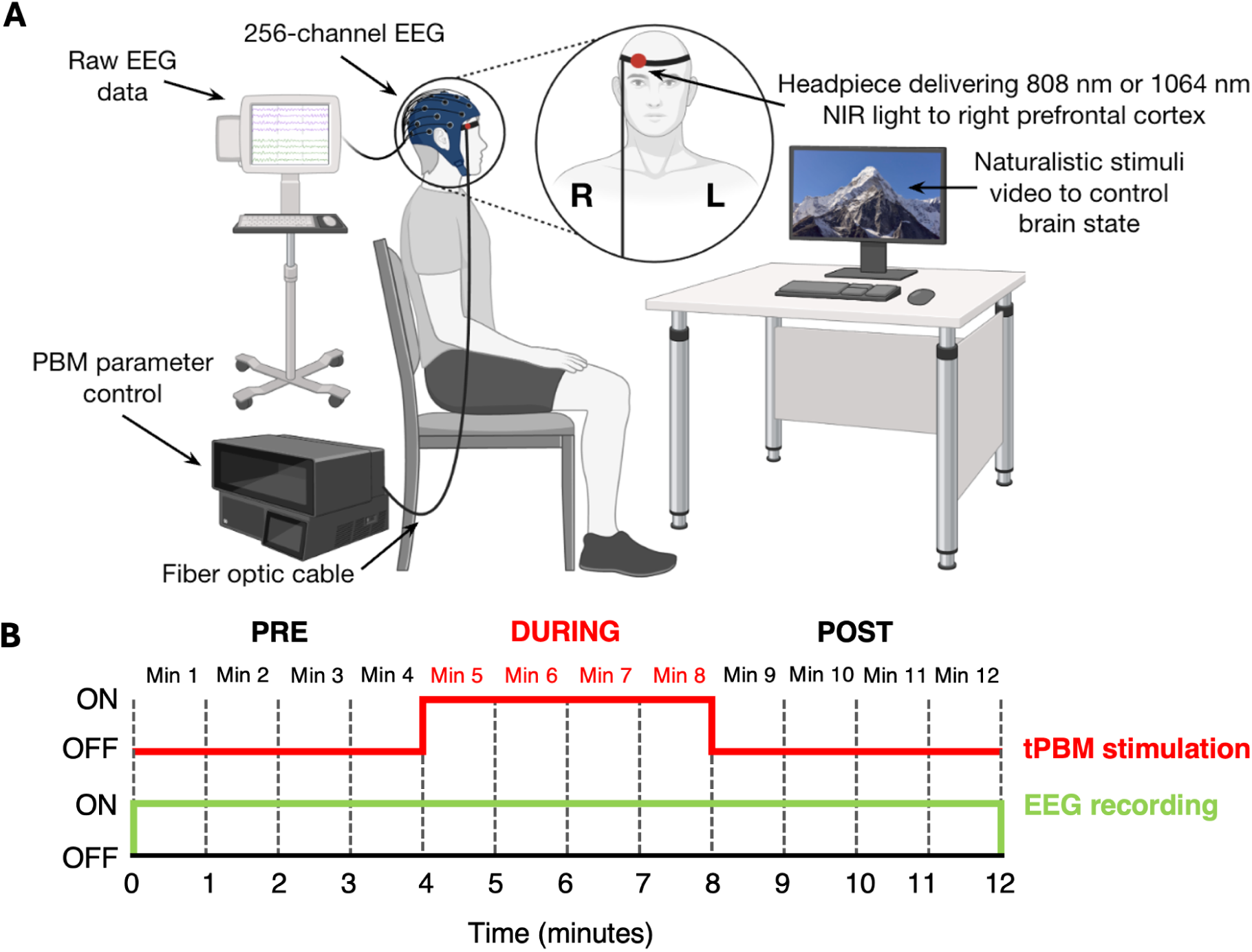
Experimental setup and recording timeline. (A) Schematic illustration of EEG-tPBM experimental setup (*BioRender.com*). (B) EEG recording timeline following block tPBM stimulus design.

**Figure 3.**
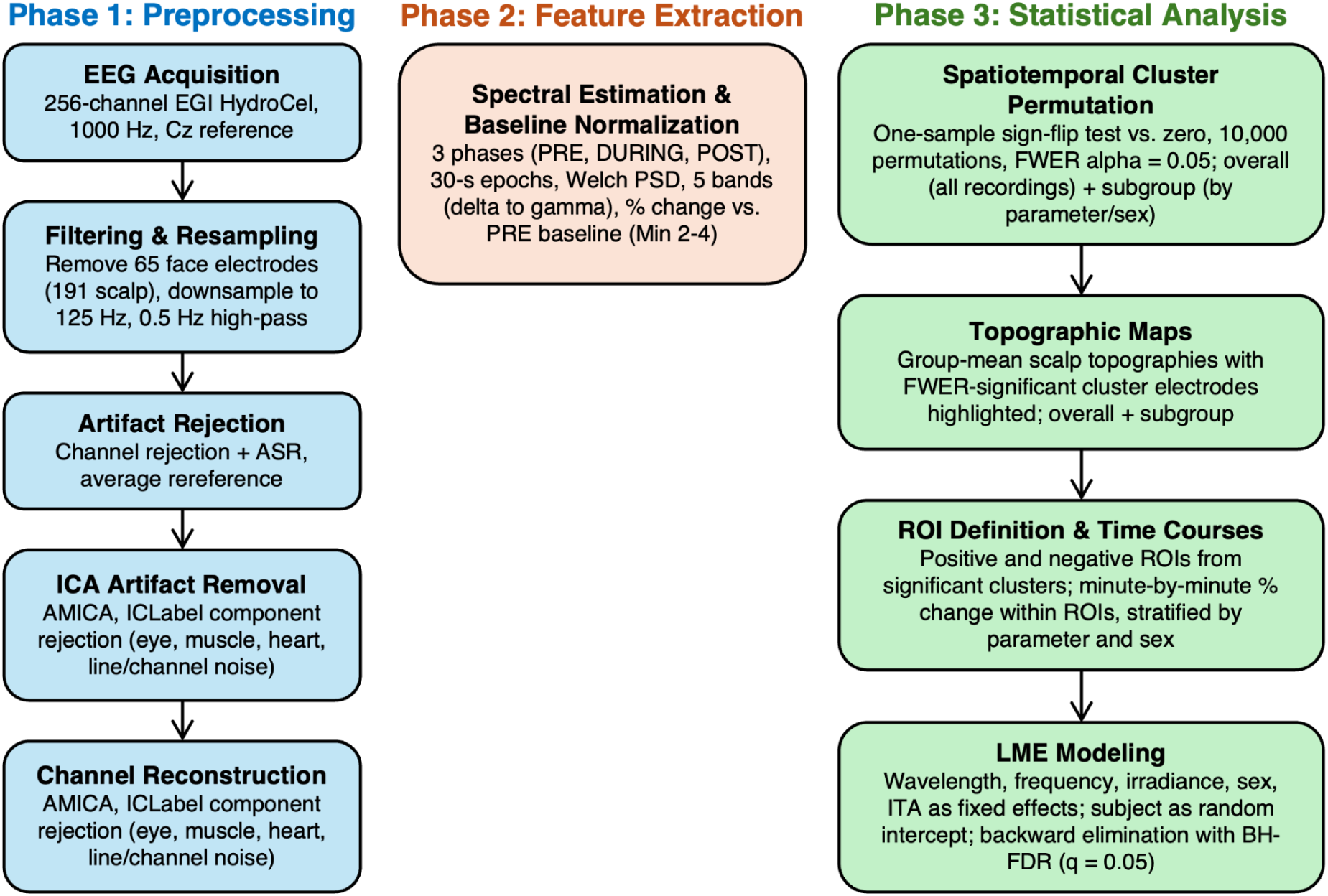
Summary of key stages in the data pipeline. EEG = electroencephalography; EGI = Electrical Geodesics Inc.; ASR = Artifact Subspace Reconstruction; ICA = Independent Component Analysis; AMICA = Adaptive Mixtures Independent Component Analysis; ICLabel = Independent Component Label; PSD = Power Spectral Density; FWER = Family-wise Error Rate; ROI = Region Of Interest; ITA = Individual Typology Angle; LME = Linear Mixed Effects; BH-FDR = Benjamini-Hochberg False Discovery Rate.

**Table 2.** Participant distribution across protocols and stimulation parameters.

| Recording | Protocol 1 | Protocol 2 | Protocol 3 |
| --- | --- | --- | --- |
| 1 | 1064 nm, 100 mW/cm<sup>2</sup>, 10 Hz | 1064 nm, 150 mW/cm<sup>2</sup>, 10 Hz | 1064 nm, 150 mW/cm<sup>2</sup>, 40 Hz |
| 2 | 808 nm, 100 mW/cm<sup>2</sup>, 10 Hz | 808 nm, 150 mW/cm<sup>2</sup>, 10 Hz | 808 nm, 150 mW/cm<sup>2</sup>, 40 Hz |
| 3 | 1064 nm, 200 mW/cm<sup>2</sup>, 10 Hz | 1064 nm, 200 mW/cm<sup>2</sup>, 40 Hz | 1064 nm, 100 mW/cm<sup>2</sup>, 40 Hz |
| 4 | 808 nm, 200 mW/cm<sup>2</sup>, 10 Hz | 808 nm, 200 mW/cm<sup>2</sup>, 40 Hz | 808 nm, 100 mW/cm<sup>2</sup>, 40 Hz |
| **M / F** | 8 M / 8 F | 7 M / 7 F | 9 M / 7 F |

**Table 3.**
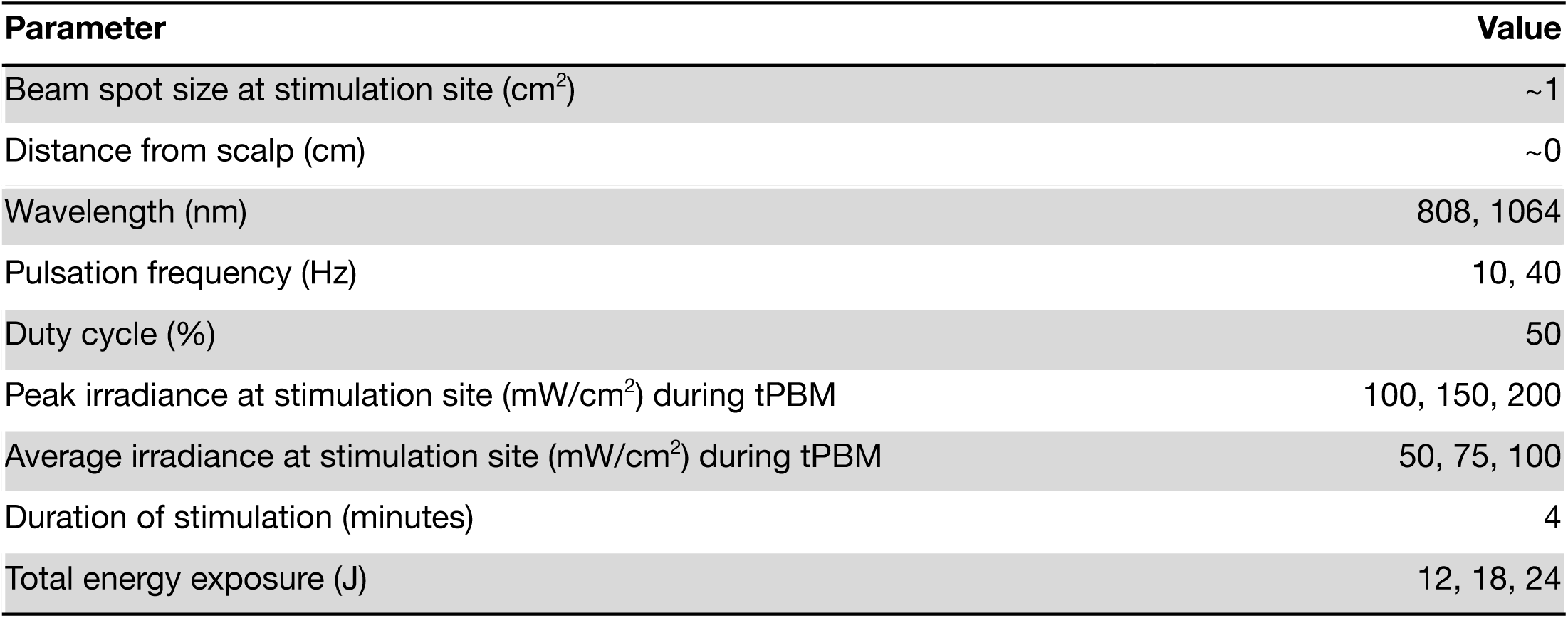
tPBM dosage parameters.

| Parameter | Value |
| --- | --- |
| Beam spot size at stimulation site (cm<sup>2</sup>) | ~1 |
| Distance from scalp (cm) | ~0 |
| Wavelength (nm) | 808, 1064 |
| Pulsation frequency (Hz) | 10, 40 |
| Duty cycle (%) | 50 |
| Peak irradiance at stimulation site (mW/cm<sup>2</sup>) during tPBM | 100, 150, 200 |
| Average irradiance at stimulation site (mW/cm<sup>2</sup>) during tPBM | 50, 75, 100 |
| Duration of stimulation (minutes) | 4 |
| Total energy exposure (J) | 12, 18, 24 |

### 2.3 EEG acquisition

EEG was recorded at 1,000 Hz using a 256-channel HydroCel Geodesic Sensor Net with a NetAmps 400 amplifier (Magstim EGI, Eugene, OR, USA; saline-based; reference Cz; impedance < 50 kΩ; amplitude resolution 0.024 µV).

### 2.4 EEG data analysis

#### 2.4.1 Preprocessing

EEG data were preprocessed in MATLAB 2025b (MathWorks Inc., Natick, MA, USA) using EEGLAB 2026.0.0.^45^ Sixty-five face electrodes were removed (191 scalp channels retained) and data were downsampled to 125 Hz. A zero-phase second-order Butterworth high-pass filter (0.5 Hz cutoff) was applied to remove slow drifts while preserving delta-band activity. Automated channel rejection and Artifact Subspace Reconstruction (ASR; channel correlation = 0.85; flat-line = 5 s; line-noise = 4; burst criterion = 10 SD) were applied via EEGLAB’s clean_rawdata; epoch rejection was disabled. An average reference was computed before Independent Component Analysis (ICA) ICA, which was performed using Adaptive Mixture ICA (AMICA)^46^ (one mixture; max 2000 iterations; PCA dimensionality = n-1). ICLabel was used to classify components; those labeled as Muscle, Heart, Line Noise, or Channel Noise (≥0.80) or Eye (≥0.70) were removed. Rejected channels were restored by spherical spline interpolation, and residual spikes exceeding twenty times the channel’s robust noise level above 300 µV were linearly interpolated.

#### 2.4.2 Frequency band extraction and power calculation

Each electrode’s data was linearly detrended over the full recording and segmented into non-overlapping 30-second epochs. Power spectral density (PSD) was estimated via Welch’s method (2-second Hann windows, 50% overlap) with sub-window periodograms combined using a 20% trimmed mean. Band-limited absolute power was integrated within predefined ranges (delta: 1-4 Hz; theta: 4-8 Hz; alpha: 8-12 Hz; beta: 12-30 Hz; gamma: 30-50 Hz) and expressed as a percent change from the mean of the PRE baseline period (Min 2-4; **Eq. 2**). Minute 1 was excluded from the baseline due to instability.

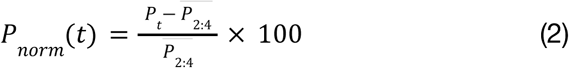

where P_t_ is the band power in minute t and 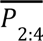 is the average band power over Min 2-4, and P_norm_ is the baseline-normalized absolute band power.

#### 2.4.3 Statistical analyses

##### 2.4.3.1 Spatiotemporal cluster-based permutation to identify significant electrode clusters

To identify changes in EEG frequency band power that were spatially and temporally consistent across subjects, cluster-based permutation tests^47^ were performed using the MNE 1.12.1 package in Python 3.11.4. To study the overall effect of tPBM as well as to observe any parameter-dependencies in the EEG responses, two group analyses were conducted. In the first analysis, all EEG recordings were averaged within-subject (**Fig. 4**) and in the second analysis, EEG recordings were first grouped by stimulation parameters or sex and then averaged within-subject (**Fig. 5-7**).

**Figure 4.**
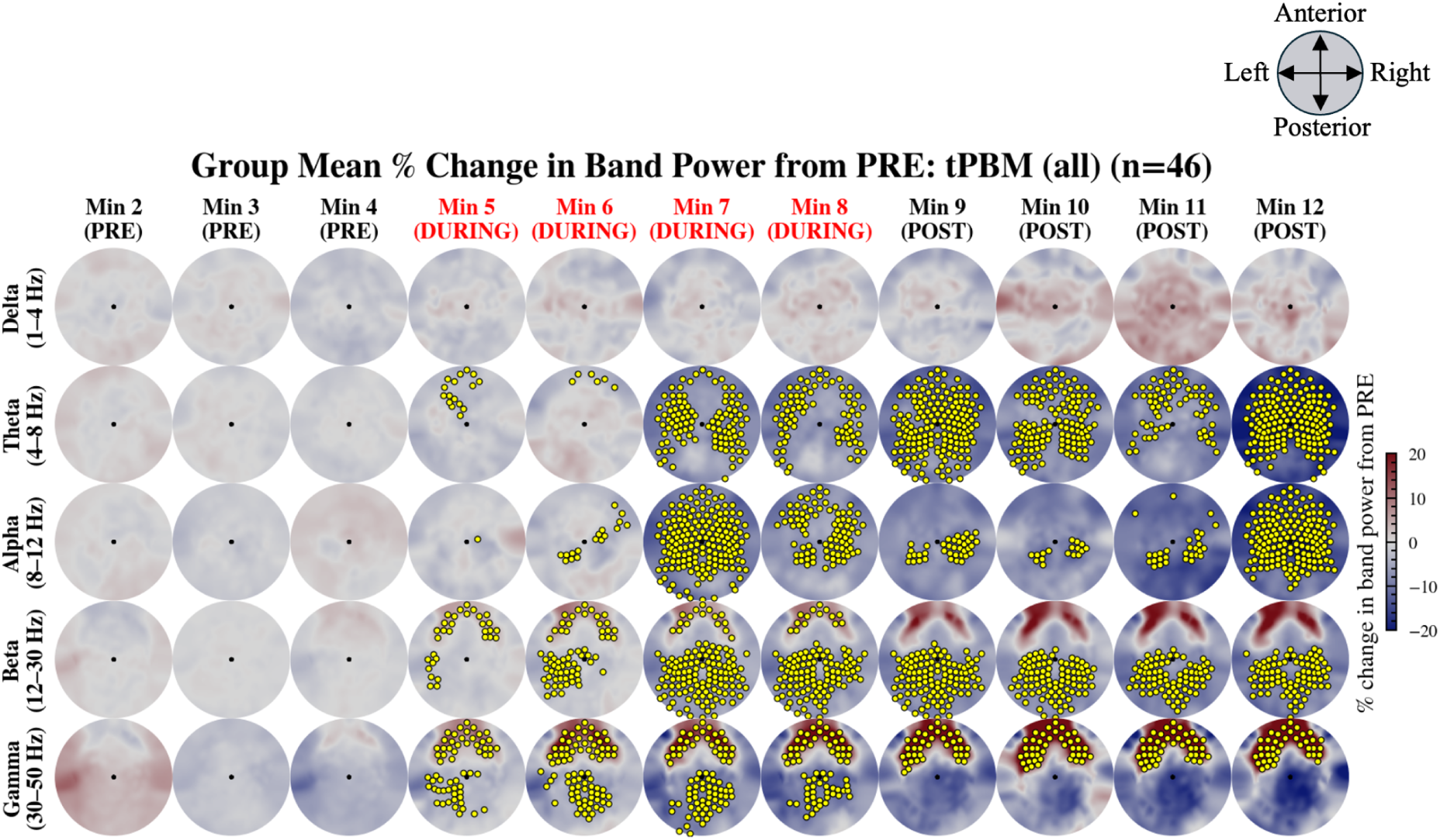
Group response to tPBM. Group-mean percent change in EEG band power from the mean of the baseline PRE period (Min 2-4) during tPBM (n=46). Rows: frequency bands (delta to gamma). Columns: successive 1-minute epochs (Min 2-12). Yellow markers: electrodes belonging to FWER-significant spatiotemporal clusters in that epoch (sign-flip permutation test, 10,000 permutations, cluster-level p<0.05). Epochs with tPBM stimulation (Min 5-8) are labeled in red. Color scale: ±20% change from PRE.

**Figure 5.**
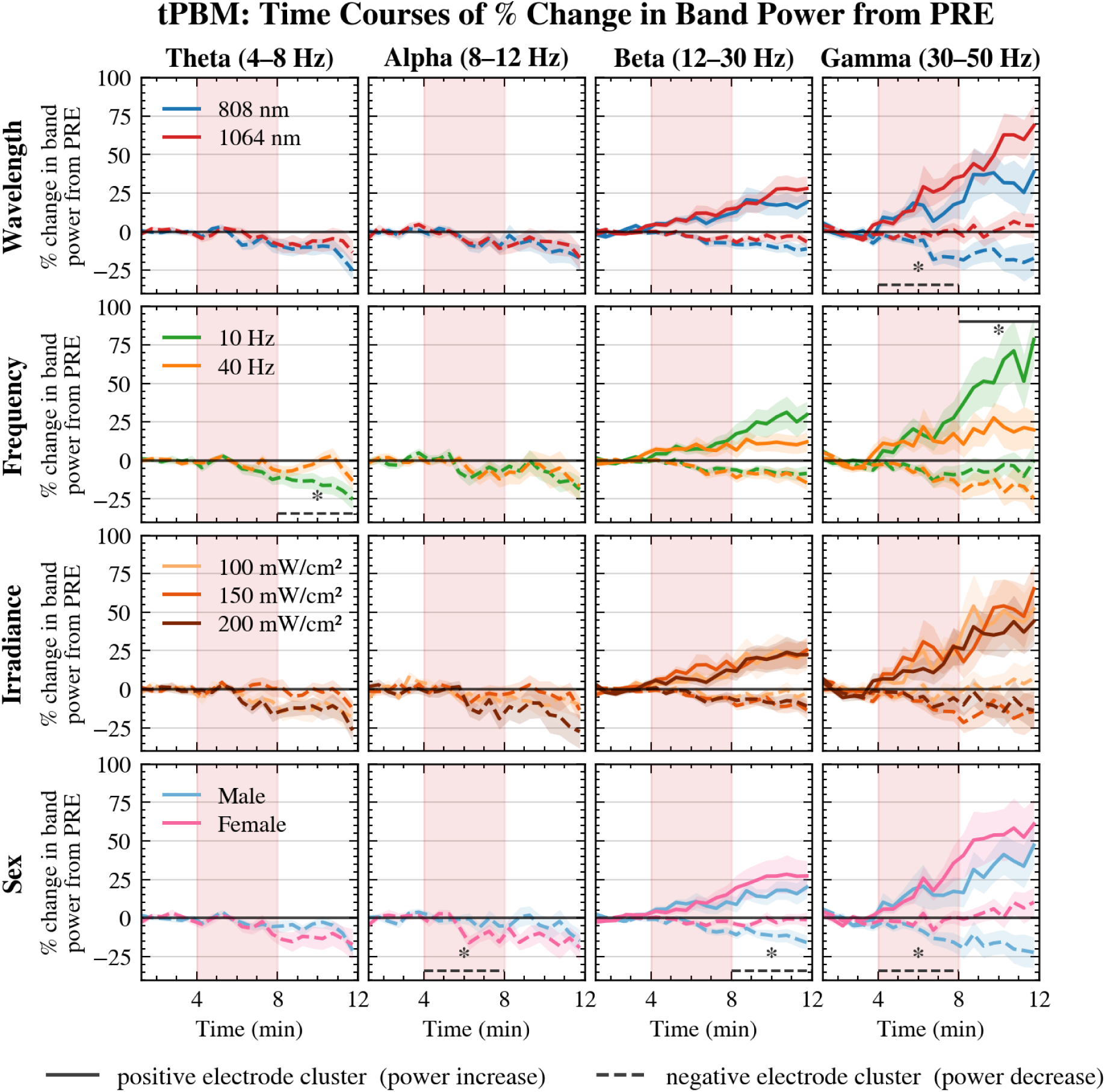
Group-level EEG band power responses to tPBM, stratified by stimulation parameters and sex. Minute-by-minute percent change in band power from the mean of the baseline PRE period (Min 2-4) is plotted within the FWER-significant positive (solid lines; power increase) and negative (dashed lines; power decrease) spatiotemporal cluster ROIs. Rows: parameter subgroups (wavelength, pulsation frequency, irradiance, sex); shaded bands indicate ±1 SEM across subjects. Columns: frequency bands with at least one significant cluster (theta to gamma). Red-shaded region: active tPBM stimulation period. Bar (solid) with asterisk: significant differences (FDR q<0.05) in parameter levels within the positive cluster ROI. Bar (dashed) with asterisk: significant differences (FDR q<0.05) in parameter levels within the negative cluster ROI. The position of the bars indicate the time window over which the significant differences were present (during or post-stimulation) (**Table 4**).

**Figure 6.**
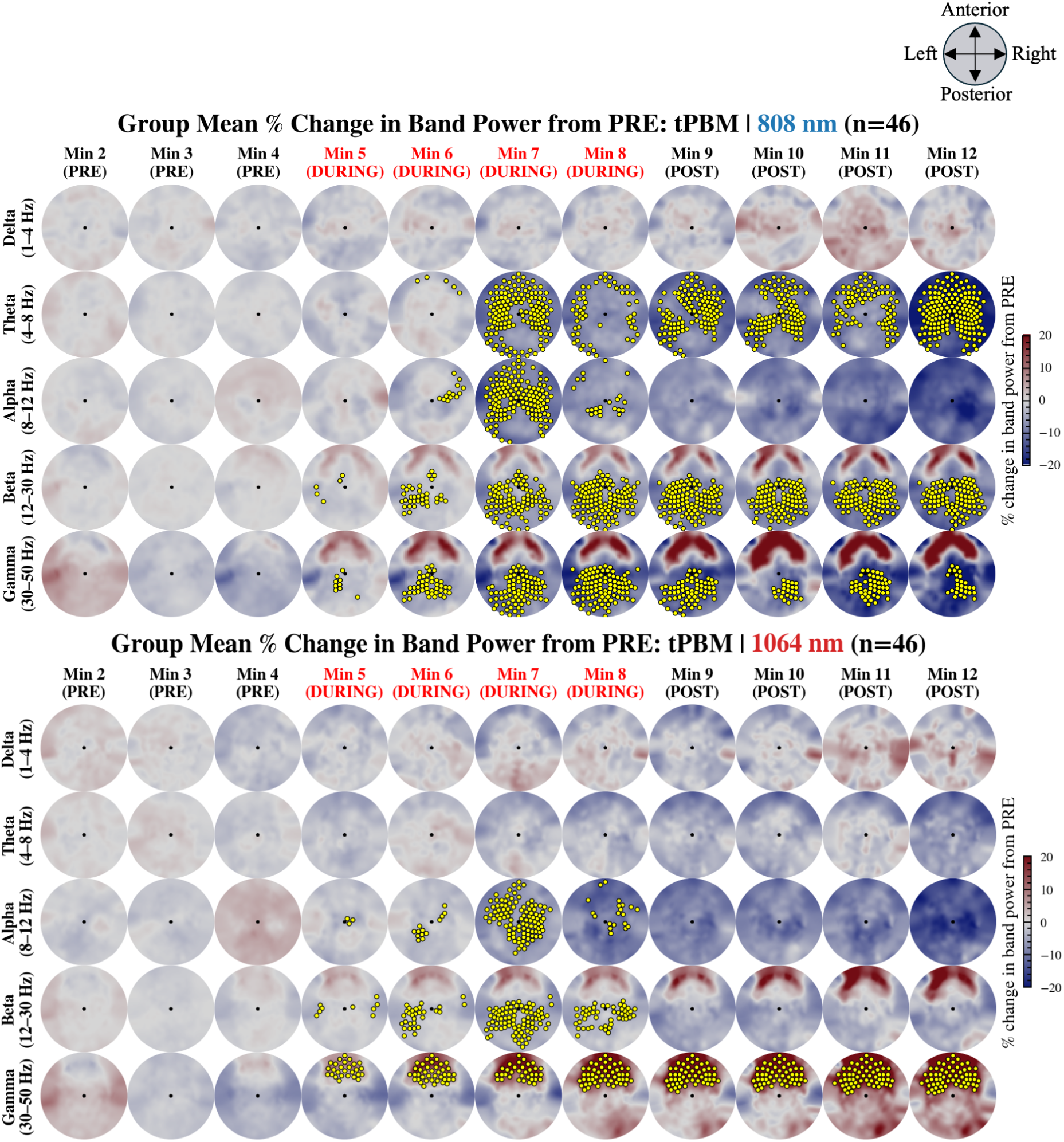
Group response to tPBM, split by wavelength. Group-mean percent change in EEG band power from the mean of the baseline PRE period (Min 2-4) during tPBM (n=46), stratified by wavelength: 808 nm (n=46) and 1064 nm (n=46). Rows: frequency bands (delta to gamma). Columns: successive 1-minute epochs (Min 2-12). Yellow markers: electrodes belonging to FWER-significant spatiotemporal clusters in that epoch (sign-flip permutation test, 10,000 permutations, cluster-level p<0.05, pooling by wavelength level. Epochs with tPBM stimulation (Min 5-8) are labeled in red. Color scale: ±20% change from PRE.

**Figure 7.**
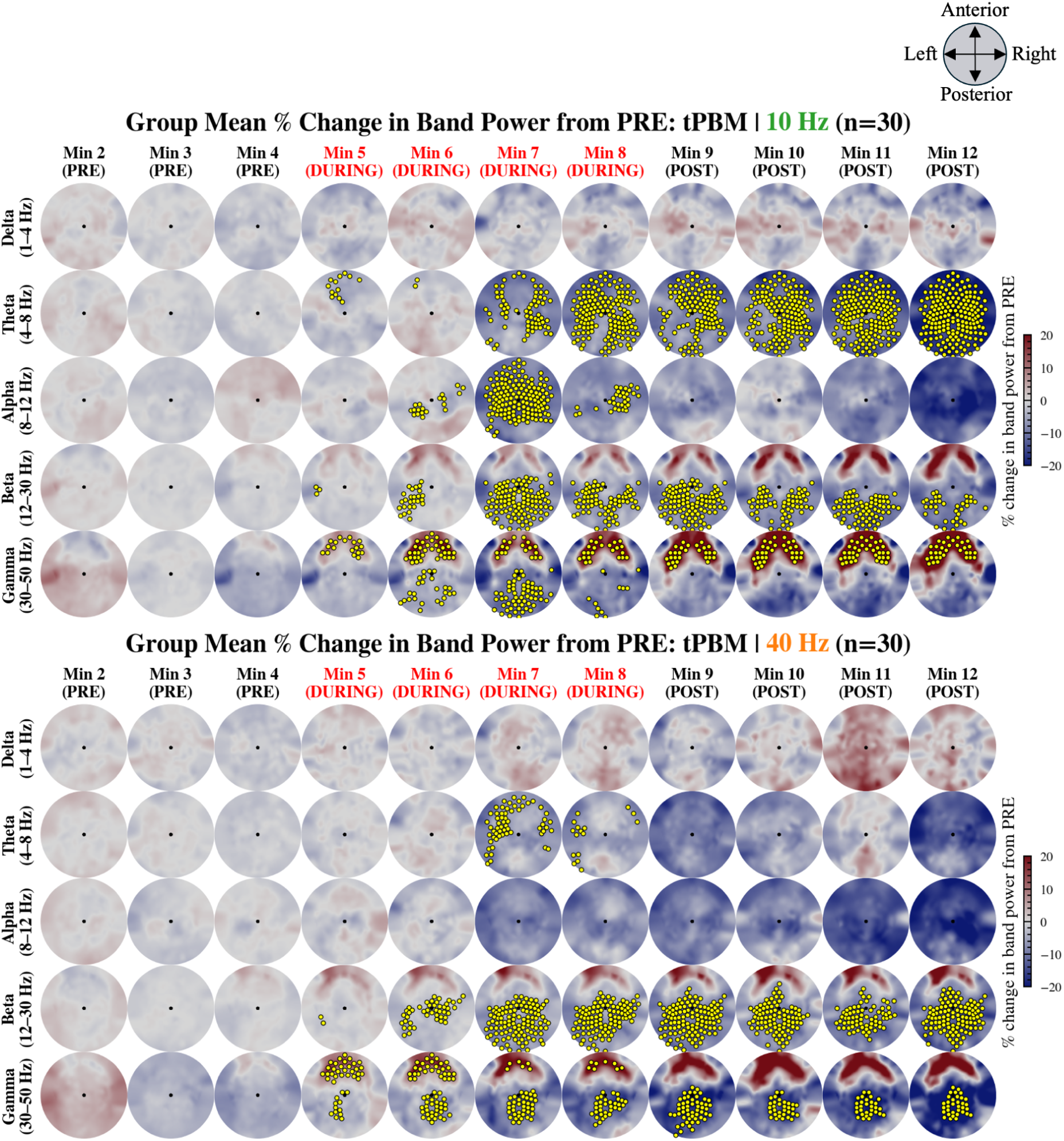
Group response to tPBM, split by pulsation frequency. Group-mean percent change in EEG band power from the mean of the baseline PRE period (Min 2-4) during tPBM (n=46), stratified by pulsation frequency: 10 Hz (n=30) and 40 Hz (n=30). Rows: frequency bands (delta to gamma). Columns: successive 1-minute epochs (Min 2-12). Yellow markers: electrodes belonging to FWER-significant spatiotemporal clusters in that epoch (sign-flip permutation test, 10,000 permutations, cluster-level p<0.05, pooling by pulsation frequency level. Epochs with tPBM stimulation (Min 5-8) are labeled in red. Color scale: ±20% change from PRE.

**Table 4.**
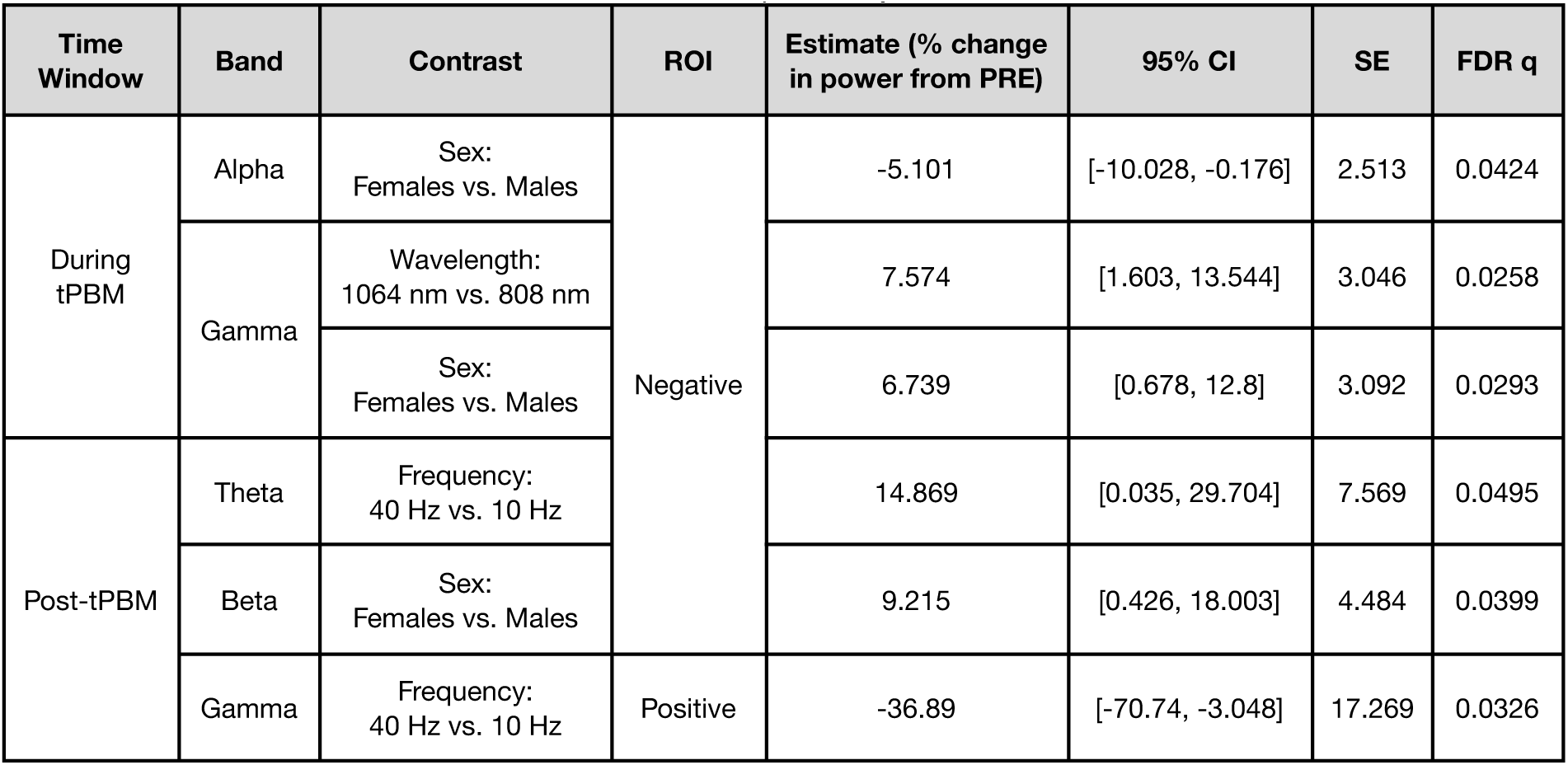
LME-derived significant parameter and biological effects. FDR-significant effects of stimulation parameters and individual biology on tPBM-induced EEG band power changes (LME model, Model A). Values are estimated differences in mean percent change from the mean of the baseline PRE period (Min 2-4) relative to the following reference levels: 808 nm wavelength, male sex, 10 Hz pulsation frequency, and 100 mW/cm^2^ irradiance. ITA is a continuous covariate; its estimate reflects the change in the EEG band power per one standard deviation increase in ITA (toward lighter skin). Full model results (non-significant contrasts and the backward elimination trace) are reported in **Tables S3 and S4**.

In each analysis, for each band x time window (DURING, POST), a 3D array (subjects ╳ time bins ╳ electrodes) was constructed where each time bin comprised 8 non-overlapping 30-second windows. Spatiotemporal adjacency was defined by combining temporal adjacency (consecutive windows) with electrode spatial adjacency (3D inter-electrode distances) via MNE’s combine_adjacency function. A one-sample sign-flip permutation test against zero was applied (two-tailed; 10,000 permutations). The cluster-forming threshold was an uncorrected alpha = 0.05 per time-electrode element. Cluster-level significance was controlled with a family-wise error rate (FWER) of alpha = 0.05 based on the maximum cluster-mass statistic across permutations.

##### 2.4.3.2 Time courses of significant electrodes

From the FWER-significant spatiotemporal clusters identified in **Section 2.4.3.1**, two regions of interest (ROIs) were defined: a positive ROI comprising electrodes belonging to any FWER-significant cluster with a positive group-mean change, and a negative ROI comprising those with a negative change. For each recording, percent-change values were averaged across ROI electrodes; recording-level time courses were then averaged within subjects; group means ±1 SEM were plotted per minute (**Fig. 8**). Only frequency bands with at least one FWER-significant cluster are shown. These same ROIs were used to visualize parameter-stratified time courses, permitting direct cross-group comparison on a common electrode set. No inferential tests were applied to time courses; parameter-level inference relied on mixed-effects models (**Section 2.4.3.3**).

**Figure 8.**
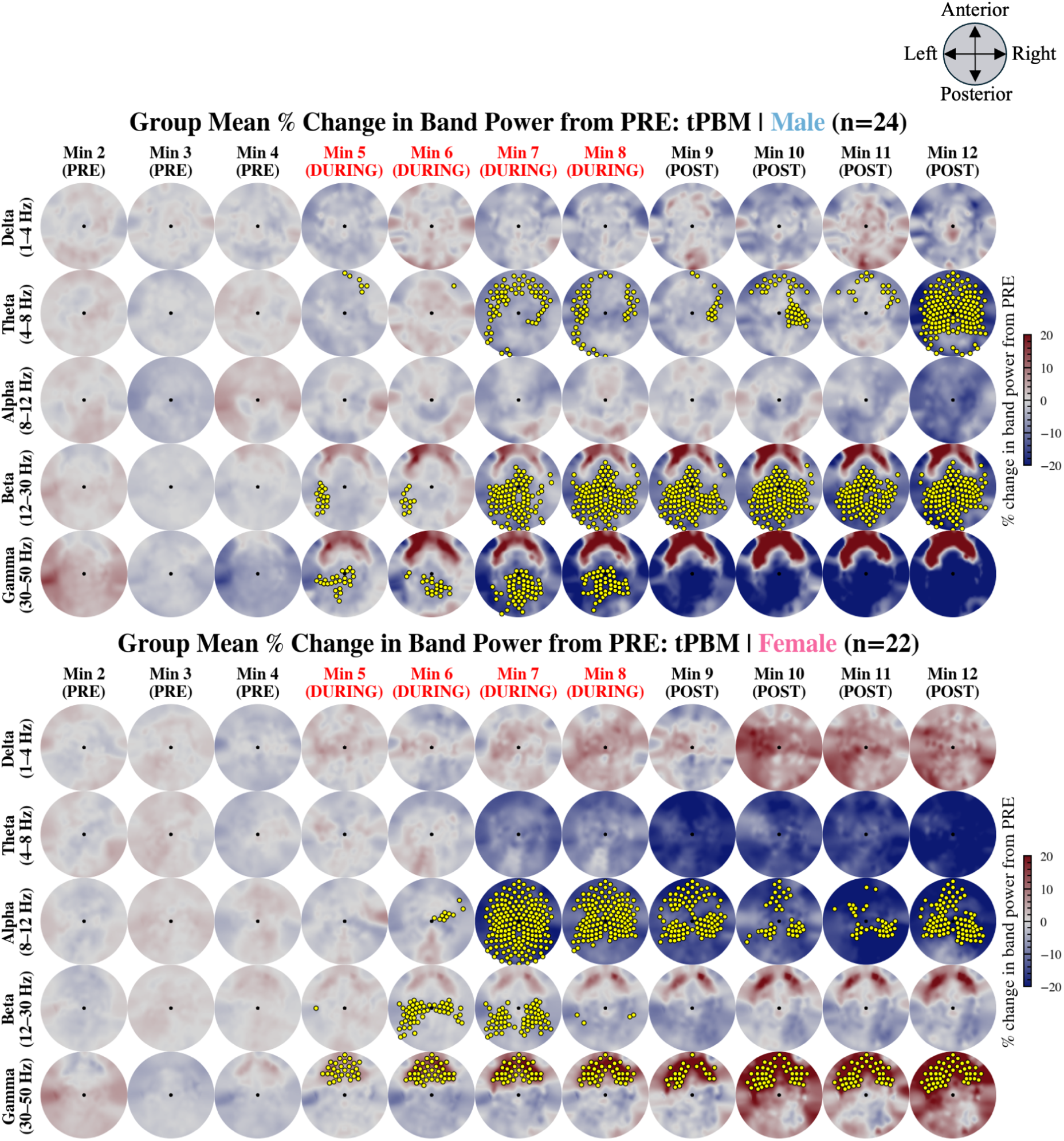
Group response to tPBM, split by sex. Group-mean percent change in EEG band power from the mean of the baseline PRE period (Min 2-4) during tPBM (n=46), stratified by sex: Male (n=24) and Female (n=22). Rows: frequency bands (delta to gamma). Columns: successive 1-minute epochs (Min 2-12). Yellow markers: electrodes belonging to FWER-significant spatiotemporal clusters in that epoch (sign-flip permutation test, 10,000 permutations, cluster-level p<0.05, pooling by sex. Epochs with tPBM stimulation (Min 5-8) are labeled in red. Color scale: ±20% change from PRE.

##### 2.4.3.3 Modeling the effects of stimulation parameters and biological factors

Linear mixed-effects (LME) models were fit in Python 3.11.5 (statsmodels v0.14; REML) to test the effects of stimulation parameters and individual biology on EEG responses within the FWER-significant electrode set. Models were fit separately for each frequency band, time window (DURING, POST), and responder class (positive ROI, negative ROI). The outcome Y was the recording-level mean percent change in band power averaged across the relevant ROI. Subject ID was included as a random intercept (experimental unit = recording session). The formula (Model A) is shown in **Eq. 3**:

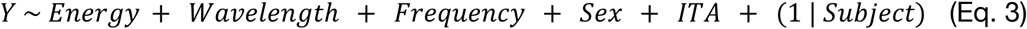

Categorical variables were Energy (Low: 100 mW/cm^2^, Mid: 150 mW/cm^2^, High: 200 mW/cm^2^), Wavelength (808 nm, 1064 nm), Frequency (10 Hz, 40 Hz), and Sex (Male, Female), while ITA was z-scored and included as a continuous variable. The reference levels were Low (Energy), 808 nm (Wavelength), 10 Hz (Frequency), and Male (Sex). Backward elimination was applied: at each step, Benjamini-Hochberg false discovery rate (FDR) correction (q=0.05) was applied simultaneously to all fixed-effect p-values, and the predictor with the highest FDR-corrected q-value was removed if it exceeded 0.05. Elimination continued until all contrasts of all remaining predictors were FDR-significant.

Model A coefficients are provided in **Table S3**, and the backward elimination trace is provided in **Table S4**. A separate carry-over analysis (Model B) tested whether the energy level of the immediately preceding recording session predicted the current response (see **Table S5**).

## 3 RESULTS

To study the spatial and temporal progression of band-specific EEG responses to pulsed forehead tPBM across key stimulation parameters (wavelength, pulsation frequency, irradiance) and biological factors (sex, skin tone), three analyses were conducted: spatiotemporal cluster-based permutation tests to identify where and when significant responses occurred (**Fig. 4-7**), time courses to describe their temporal evolution within significant electrode regions (**Fig. 8**), and LME modeling to quantify the contribution of each parameter and biological factor to the response magnitude (**Table 4**). Notably, the carry-over analysis revealed that no tPBM-EEG recordings were significantly affected by the energy level of the previous recordings in our study (see **Table S5** for more details).

### 3.1 Spatiotemporal EEG response to pulsed forehead tPBM

**Fig. 4** shows the group-mean scalp topographies for all 46 participants across all recorded minutes. No significant spatiotemporal clusters were identified for delta power. Theta power exhibited significant frontal decreases beginning at the onset of stimulation (Min 5) that grew stronger and became more widespread and more significant in the second half of the stimulation period (Min 7), continuing through the post-stimulation period. Alpha power similarly decreased from baseline, with significant clusters emerging during the final two minutes of stimulation (Min 7-8) and the largest and most spatially extensive reductions observed in the final post-stimulation minutes, resembling the theta power reductions. Beta and gamma power responses were bidirectional: both bands displayed significant posterior reductions and anterior increases during stimulation, however, post-stimulation, posterior reductions only remained in the beta band and anterior increases only remained in the gamma band.

### 3.2 Parameter and biological dependencies in the spatiotemporal EEG response to pulsed forehead tPBM

**Fig. 5** plots minute-by-minute percent changes in EEG band power from the pre-stimulus baseline, stratified by stimulation parameters and sex. At the low-frequency end of the spectrum, delta power was not significantly modulated by any parameter or sex. Theta power, however, exhibited reductions across all parameter levels and sexes but only pulsation frequency significantly modulated the magnitude of this effect post-stimulation, with 40 Hz associated with smaller reductions than 10 Hz (by 14.9%, 95% CI [0%, 29.7%]) (**Table 4**). In the mid-frequency range, alpha power decreased from baseline across all parameter levels, with sex emerging as the only significant modulator during stimulation: females displayed larger reductions than males (by 5.1%, 95% CI [0.2%, 10%]) (**Table 4**). At the high-frequency end, beta and gamma responses were bidirectional and showed prominent parameter- and sex-level separation. For beta, only sex significantly modulated the post-stimulation response: females displayed smaller reductions than males (by 9.2%, 95% CI [0.4%, 18%]) (**Table 4**). For gamma, wavelength and sex were significant modulators during stimulation: 1064 nm was associated with smaller reductions than 808 nm (by 7.6%, 95% CI [1.6%, 13.5%]), and females showed smaller gamma reductions than males (by 6.7%, 95% CI [0.7%, 12.8%]) (**Table 4**). In the post-stimulation period, pulsation frequency significantly modulated the positive gamma response, with 40 Hz producing smaller increases than 10 Hz (by 36.9%, 95% CI [3%, 70.7%]) (**Table 4**).

Based on the significant parameter effects identified by the LME, the spatiotemporal profiles of the EEG responses were further stratified by wavelength, pulsation frequency, and sex (**Fig. 6-8**).

**Fig. 6** compares the scalp topographies between 808 nm (n=46) and 1064 nm (n=46) pulsed forehead tPBM. Both wavelengths produced broadly similar spatial patterns, however, 808 nm resulted in more spatially extensive and temporally persistent FWER-significant electrode clusters across theta, beta, and gamma bands. Neither wavelength produced significant delta modulation. For 808 nm, theta power was significantly reduced in a widespread pattern beginning in the second half of stimulation (Min 7) and persisting throughout the post-stimulation period, with the strongest effects at the end of the recording. Alpha reductions reached significance in the second half of stimulation. Beta and gamma power were both significantly reduced in posterior regions from stimulation onset, with effects growing in magnitude through the end of the post-stimulation period – gamma showing stronger changes than beta. For 1064 nm, no significant theta modulation was identified. Alpha reductions were more prominent and spatially widespread than for 808 nm, emerging in the second half of stimulation. Beta power was significantly reduced posteriorly but only during stimulation. Gamma power, by contrast, showed a significant frontal and prefrontal increase during and post-stimulation that grew progressively in magnitude across the recording.

**Fig. 7** compares the scalp topographies between forehead tPBM pulsed at 10 Hz (n=30) and 40 Hz (n=30). Neither frequency produced significant delta modulation. With 10 Hz, theta power was significantly reduced beginning in the second half of stimulation, becoming more widespread as stimulation progressed and into the post-stimulation period. Alpha reductions were significant during the second half of stimulation but did not persist post-stimulation. Beta power was significantly reduced in posterior regions both during and post-stimulation. Gamma power showed significant frontal and prefrontal increases during and post-stimulation, growing across the recording, while significant gamma reductions were limited to the stimulation period. For 40 Hz, no significant alpha modulation was identified. Theta reductions were significant in the second half of stimulation but spatially limited to frontal and parietal regions. Beta power was significantly reduced posteriorly both during and post-stimulation. Gamma showed significant frontal and prefrontal increases during stimulation. Significant gamma reductions, however, were present both during and post-stimulation and grew stronger into the post-stimulation period — a pattern distinct from the 10 Hz condition.

**Fig. 8** compares the scalp topographies between the male (n=24) and female (n=22) response to pulsed forehead tPBM. Neither group showed significant delta modulation. In males, theta power was significantly reduced in the second half of stimulation and became more widespread only at the very end of the recording (Min 12). No significant alpha modulation was observed in males. Beta power was significantly reduced in posterior regions both during and post-stimulation, growing stronger post-stimulation. Gamma power was significantly reduced posteriorly during stimulation. In females, no significant theta modulation was observed. Alpha power was significantly and widely reduced in the second half of stimulation (Min 7), with the spatial extent of significant clusters decreasing as the recording progressed. Beta reductions were significant and confined to posterior regions only during stimulation. Gamma power showed significant frontal and prefrontal increases both during and post-stimulation, with the effect growing in magnitude into the post-stimulation period – a response pattern not seen in males.

## 4 DISCUSSION

In 46 healthy young adults, pulsed forehead tPBM robustly modulated resting-state EEG power in a frequency band- and parameter-dependent manner, contributing to varying spatiotemporal profiles of online (during stimulation) and offline (post-stimulation) effects. Our findings reveal that tPBM (i) evokes bilateral EEG responses that were not time-locked to the block stimulus shape, persisting post-stimulation; (ii) significantly increases anterior high-frequency (gamma) power while significantly reducing posterior low-to-mid-frequency (theta to beta) power; (iii) typically affects beta and gamma power earlier than it does theta and alpha; (iv) effects were only significantly modulated by differences in wavelength, pulsation frequency, and participant sex.

### 4.1 Pulsed forehead tPBM evokes bilateral EEG responses not time-locked to stimulation

Despite only irradiating a 1 cm^2^ site on the right forehead, tPBM produced bilateral responses (**Fig. 4**), consistent with fNIRS studies, which measure changes in the concentrations of oxygenated (HbO) and deoxygenated hemoglobin (dHbO), the downstream hemodynamic consequences of altered neural metabolism. Because tPBM is proposed to enhance oxidative phosphorylation and release NO, which acts as a vasodilator, fNIRS provides a particularly direct window into whether the metabolic cascade initiated by CCO stimulation translates into measurable hemodynamic change. Indeed, bilateral prefrontal hemodynamic responses from unilateral forehead stimulation have been repeatedly documented: Tian et al. found simultaneous bilateral HbO increases and deoxy-HbO decreases,^14^ Wang et al. observed bilateral changes in metabolic oscillations and prefrontal functional connectivity,^48^ and Saucedo et al. and Urquart et al. reported similar bilateral patterns.^17,20^ Zomorrodi et al. also reported bilateral EEG changes following pulsed combined tPBM.^29^ The convergence of EEG and fNIRS on the bilateral spatial nature of responses to tPBM, with one measuring cortical oscillatory activity and the other the hemodynamic consequences of mitochondrial engagement, points to a neural response that manifests at different timescales.

The right prefrontal cortex is a hub within bilateral resting-state networks, including the frontoparietal network (FPN),^49,50^ and locally enhanced metabolism likely propagates through this existing connectivity to recruit distributed bilateral nodes. Urquhart et al. directly observed this propagation: functional connectivity was most significantly modulated in the FPN and regions of other brain networks that are highly integrated with the FPN, suggesting a potential network-level reorganization that outlasted stimulation.^17^ That both 808 nm and 1064 nm produced this same widespread signature, despite partly distinct biological mechanisms (**Section 4.3**), further supports network propagation as the common pathway.

Effects also emerged gradually from the second half of stimulation and continued growing post-stimulation (**Fig. 4-8**). This temporal profile closely mirrors the fNIRS literature: immediate rises in oxidized CCO during stimulation – the primary photobiological event – followed by a temporally offset HbO increase and dHbO decrease that peaked only after light offset,^20^ demonstrating that the metabolic trigger and its hemodynamic consequence are separated by minutes. This ramp-like persistence could be reflective of the timescale of the photobiological cascade, including mitochondrial redox changes, ATP synthesis, NO-mediated vasodilation, and ion-channel modulation, which may operate over minutes, rather than milliseconds. The EEG changes we measure may reflect the end-readout of this cascade, not a direct reflection of photon delivery, implying that the post-stimulation period should be considered an integral part of the response rather than merely a return-to-baseline control.

### 4.2 Pulsed forehead tPBM increases anterior high-frequency EEG power and reduces posterior low-to-mid-frequency power

The response was a spatially organized spectral reorganization: frontal gamma enhancement, consistent with Zomorrodi et al.^29^ and Spera et al.^51^, alongside posterior theta, alpha, and beta suppression. This dual pattern suggests two concurrent phenomena: local cortical enhancement in the stimulated prefrontal tissue (gamma) and distributed arousal propagating through connected networks (posterior suppression). Our alpha suppression contrasts with previous reports of alpha increases,^52,53^ likely reflecting our use of pulsed rather than continuous wave delivery and the eyes-open, video-engaged protocol. Beta and gamma responded earliest (from stimulation onset), while theta and alpha emerged later (Min 7-8), consistent with their distinct neural generators: fast-spiking interneuron-dependent high-frequency rhythms respond quickly to local metabolic changes, whereas slower rhythms require longer network-level reorganization.^54^

### 4.3 Parameter choice matters: the significant effect of wavelength and pulsation frequency

Wavelength qualitatively shifted the spectral profile rather than merely scaling its magnitude. At 808 nm, the dominant response was broadband posterior suppression across theta, beta, and gamma; at 1064 nm, this suppressive signature was largely absent, replaced by progressive frontal gamma enhancement (**Fig. 5**; **Table 4**). This dissociation maps onto the distinct biological targets of each wavelength: 808 nm sits near CCO’s absorption peak^39,40^ and drives the metabolic cascade – NO photodissociation, enhanced ATP synthesis, and downstream signaling – whose network-level effects is widespread posterior suppression propagating through connected resting-state networks (**Section 4.1**). At 1064 nm, CCO absorption is weaker but penetration depth is greater and interaction with water is stronger,^33,55^ shifting the mechanism toward thermally sensitive pathways such as TRP channel gating and membrane capacitance changes that produce a more localized, frontal excitability increase.^35–37^ This is, to our knowledge, the first EEG evidence that wavelength determines the character, not merely the magnitude, of the cortical response to tPBM, and it underscores the importance of treating wavelength as a mechanistically meaningful parameter.

#### 4.3.1 Pulsed tPBM does not entrain EEG rhythms to the stimulation frequency

A key objective of this work was to test whether pulsed NIR light entrains cortical rhythms toward the stimulation frequency, as predicted by the entrainment framework for periodic stimuli.^21,56,57^ We hypothesized that 10 Hz and 40 Hz pulsing would preferentially alter alpha and gamma activity, respectively, an expectation supported by preclinical work linking 10 Hz pulsing to robust mitochondrial responses^58^ and by the prominence of 40 Hz in gamma-focused human paradigms.^27,29^ Our results directly contradict this prediction: 10 Hz pulsation produced significantly larger post-stimulation gamma increases and theta reductions than 40 Hz (**Table 4**), meaning that the lower, not the higher, pulsation rate drove the larger high-frequency response.

This pattern parallels the sensory-stimulation literature. Just as 40 Hz flickering light evokes only a narrow-band steady-state response without engaging endogenous gamma,^28^ pulsed tPBM at 40 Hz did not preferentially amplify gamma oscillations. The pulsation frequency dependence we observe is therefore more likely photobiological than oscillatory: at a 50% duty cycle, 10 Hz delivers 50 ms continuous on-intervals per cycle versus 12.5 ms at 40 Hz, potentially allowing greater cumulative CCO photostimulation per cycle before the enzyme transitions back to its ground redox state.^58^

#### 4.3.2 The absence of significant irradiance effects

Across our tested range (100-200 mW/cm^2^), no irradiance-dependent differences survived FDR correction in any band or time window. Parameter-stratified time courses overlapped at the group level with no consistent trend with increasing irradiance (**Fig. 8**). PBM dose-response relationships are frequently nonlinear and at times biphasic at the cellular scale, with intermediate energy levels being optimal and higher energy levels yielding diminishing returns.^59,60^ Our tested irradiances may therefore sit at or beyond the plateau of the dose-response curve. A null result is not proof of no effect; per-level sample sizes (∼15 subjects) may have been insufficient to detect small dose-dependent differences however, these results argue against assuming that the EEG response scales proportionally with delivered energy within clinically relevant irradiances, and suggest that protocol optimization is better focused on wavelength and pulsation frequency selection.

### 4.4 The importance of individual biology: the significant effect of sex, but not skin tone

Sex was the most prominent biological modulator of the tPBM EEG response, with FDR-significant effects across three band-time window combinations: females showed larger alpha suppression during stimulation, smaller posterior gamma reductions during stimulation, and smaller posterior beta reductions post-stimulation (**Table 4**, **Fig. 7**). These represent qualitatively distinct spectral profiles rather than mere magnitude differences; the female response was characterized by stronger mid-frequency cortical desynchronization (alpha) and relatively preserved posterior high-frequency power, while the male response was characterized by deeper and more sustained posterior beta and gamma suppression.

These sex-dependent tPBM responses are likely rooted, at least in part, in pre-existing sex differences in baseline high-frequency EEG power.^61,62^ Sawicka et al. recently demonstrated in a comparable cohort of healthy young adults that between-sex differences in resting-state spectral power are widespread and predominantly concentrated in beta and gamma bands over frontal and central regions, with males exhibiting significantly higher power than females.^63^ Critically, these differences were more prominent than within-sex hormonal fluctuations across the female menstrual cycle, and no within-male testosterone-EEG correlations survived FDR correction despite a substantial inter-individual range, indicating that the sex differences reflect long-term organizational shaping of cortical architecture rather than acute modulation by circulating hormone levels.^63^ If males and females begin each tPBM session from different baseline positions in the bands where we observe sex-dependent responses (alpha, beta, gamma), then the same photobiological perturbation would naturally produce different magnitudes of change, even if the underlying cellular mechanism is identical. Sex hormones may further contribute through acute receptor-mediated modulation of excitation-inhibition balance: estrogen increases neuronal excitability while progesterone shifts the balance toward inhibition via GABA-A receptor modulation,^64,65^ and estradiol has been specifically associated with posterior high gamma power,^63^ the same band and region where we observe sex-dependent tPBM effects. Because we did not collect hormone-level data, we cannot distinguish hormonal from anatomical contributors in the present dataset, and future tPBM studies should include such measures.

ITA (skin tone), on the other hand, included as a continuous proxy for melanin-related optical absorption at the forehead, showed no significant association with the EEG response in any band or time window. Because melanin strongly absorbs NIR light, participants with darker skin tones were hypothesized to experience attenuated energy reaching the cortex.^33,66^ The null finding, however, should be interpreted cautiously; our cohort may have lacked the range of skin tones or statistical power to resolve what could be a modest attenuation effect. A more definitive test will require larger, more representative samples, ideally paired with concurrent NIRS to independently quantify cortical dose at each ITA level.^33^

### 4.5 Potential mechanisms: from photon absorption to oscillatory reorganization

The band-specific, persistent, wavelength-dependent, and irradiance-insensitive EEG responses may be best explained by a multistage cascade. CCO photostimulation enhances oxidative phosphorylation,^14,20^ however EEG changes did not scale with irradiance, inconsistent with a purely energy-driven account. More likely, mitochondrial activation sets a permissive metabolic state that alters excitation-inhibition balance through redox signaling, calcium-linked pathways,^67^ and downstream ion-channel modulation,^7^ producing frequency-specific redistribution rather than uniform firing increases.^54^ The wavelength dissociation further suggests a mixture of CCO-mediated (metabolic) and water-mediated (thermal-excitability) routes.

### 4.6 Limitations and future work

Despite the foundation provided by our work, several limitations constrain the generalizability of our results. Our sample comprised healthy young adults, therefore, EEG responses may differ in patient populations where mitochondrial dysfunction is more prevalent. Our repeated-recording design, while enabling within-participant parameter comparisons, may have introduced carry-over effects from prior doses beyond what was eliminated by Model B, therefore, future work would benefit from greater temporal separation or counterbalanced ordering between stimulation conditions. Furthermore, the fixed recording order also means that fatigue or habituation effects cannot be fully disentangled from parameter effects beyond what Model B screened. Additionally, because our primary aim was to characterize parameter-specific effects across a multidimensional stimulation space, incorporating a dedicated sham condition for every parameter combination was not feasible; instead, the within-subject PRE baseline served as the reference, which is sufficient for detecting within-session changes but cannot explicitly rule out expectation effects. Our 12-minute recording window may not have fully captured the duration of post-stimulation effects, particularly given that several bands were still showing growing changes at the final time point. Our study was also limited to a single stimulation site, the right prefrontal cortex, and we did not collect concurrent behavioral or cognitive measures, so the oscillatory changes we observe cannot be directly linked to functional outcomes, though the spectral profile aligns with patterns associated with improved attention in the broader EEG literature. Given the significant sex differences identified, future studies may benefit from collecting blood markers and hormone-level data to disentangle hormonal from anatomical contributors. Finally, since EEG indexes tPBM effects only at the level of scalp cortical activity, it cannot reveal whether stimulation also reaches subcortical structures that are frequently compromised in neurodegeneration.

These limitations define a clear agenda for future work. The unexpected inverse relationship between pulsation frequency and gamma response magnitude calls for a systematic exploration of a wider range of pulsation frequencies and duty cycles, ideally paired with concurrent mitochondrial readouts such as broadband NIRS. The wavelength-dependent spectral dissociation we report (808 nm driving suppression, 1064 nm driving frontal gamma enhancement) was not tested as a formal interaction with pulsation frequency, a contrast that could further elucidate whether CCO-mediated and water-mediated mechanisms contribute differently to the pulsation frequency effect, and one that can be addressed with the existing dataset. Larger, more diverse cohorts are required to determine whether the irradiance and skin-tone null findings reflect true dose insensitivity or insufficient statistical power. Whether alternative delivery routes that bypass scalp optical properties, such as intranasal PBM, can achieve more efficient cortical modulation per unit energy remains an open and clinically important question, a direction we intend to pursue. Finally, integrating concurrent hemodynamic imaging with EEG would allow direct testing of the metabolic-to-excitability cascade we propose, rather than inferring it from temporal parallels alone.

### 4.7 Conclusions

Pulsed forehead tPBM reliably altered resting-state EEG in healthy young adults, producing posterior low-to-mid-frequency suppression and frontal gamma enhancement that developed gradually and persisted after stimulation. These effects were parameter-dependent: 808 nm drove broadband suppression while 1064 nm selectively enhanced frontal gamma, and 10 Hz pulsation produced larger effects than 40 Hz, contradicting entrainment predictions. Sex was the most consistent biological modulator; irradiance and skin tone were not significant within the tested range. These findings highlight the importance of prioritizing wavelength, pulsation frequency, and sex in tPBM protocol design, and that the post-stimulation window is also an integral component of the response profile.

## 5 DATA AVAILABILITY

Data and software code can be made available upon request.

