## Supplementary Materials for "Pulsed transcranial photobiomodulation evokes sustained, parameter-dependent changes in EEG oscillations that run opposite to frequency entrainment"

#### S1 Skin tone measurement

##### S1.1 Individual typology angle (ITA)

Skin tone was quantified using the individual typology angle (ITA), calculated using **Eq. 1**:

$$ITA = arctan(\frac{L^{*}-50}{b^{*}}) \times \frac{180}{\pi}. \quad (1)$$

where L* represents luminance ranging from black (0) to white (100), and b* represents the blue-yellow chromaticity axis. Higher ITA values correspond to lighter skin. ITA skin color types are typically classified into six ranges: very light (>55°), light (41°-55°), intermediate (28°-41°), tan (10°-28°), brown (-30°-10°), and dark (<-30°).

##### S1.2 Spectrophotometer calibration and measurement

Prior to data collection for each participant, the spectrophotometer was calibrated using both black (zero) and white reference standards. For zero calibration, the spectrophotometer was held upward with no visible objects within 1 metre of the measuring window, and a measurement was taken of the air. White calibration was performed subsequently, with a white cap attached to the end of the spectrophotometer for a second measurement.

Following calibration, the spectrophotometer was positioned over the right forehead, where the laser light would be delivered, providing a direct estimate of the skin tone at the stimulation site. Three consecutive measurements were taken and averaged for each calculation, and two rounds were used to calculate the ITA, which was then recorded per participant.

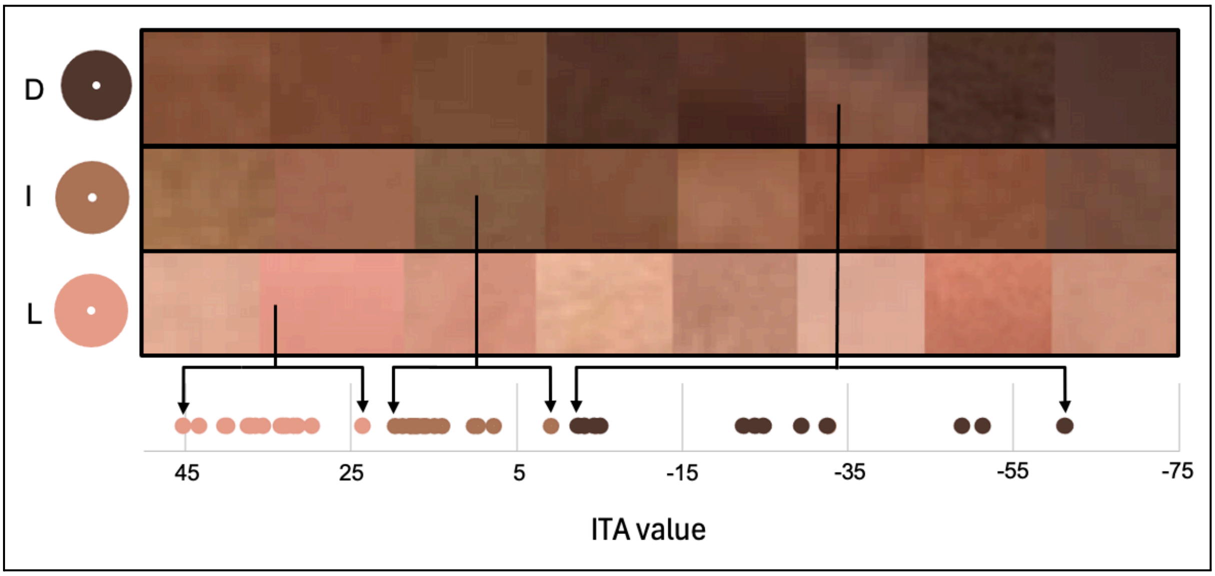


**Figure S1. Objective measurements of skin tone compared to photographs of forehead skin for a representative group of subjects.** Images were categorized based on their Individual Typology Angle (ITA) to show the general tone of each skin tone group: light (L), intermediate (I), and dark (D). Dots represent the actual ITA values corresponding to the mean visually perceived skin tone based on individual photographs of skin patches, ordered by ITA within each group. Higher ITA values reflect lighter skin.

### S2 Stimulation calibration and dosimetry

#### S2.1 Optical power calibration

Prior to data acquisition, laser output was measured and calibrated using a Newport optical power meter (Model 843-R, Newport Corporation, USA). The optical fibre was connected to the laser power supply on one end and the forehead applicator on the other. The applicator was placed directly on the power meter sensor to ensure that the measured power reflected the delivered output to the skin surface.

The signal generator was connected to the power supply using a BNC cable. The laser mode was set to modulation, and the signal generator was configured to the target pulsation frequency (10 Hz or 40 Hz) with a 50% duty cycle and 5V amplitude.

#### S2.2 Energy and irradiance dose calculation

The applied irradiance was calculated by dividing the measured optical power (mW) by the cross-sectional area of the beam at the applicator tip, reflecting the time-averaged power across pulse cycles. The beam diameter was approximately 11.28 mm, corresponding to a radius of 0.565 cm and an overall area of 1cm$^2$ ($A = \pi * r^2$). Delivered energy (J/cm$^2$) was calculated as the product of applied irradiance (mW/cm$^2$), duty cycle and stimulation duration (240 s).

The applied irradiance was calculated by dividing the measured time-averaged optical power (mW) by the cross-sectional area of the beam at the applicator tip. The beam diameter was approximately 11.28 mm, corresponding to a radius of 0.565 cm and an area of approximately 1 cm$^2$ ($A = \pi * r^2$). Delivered energy density (J/cm$^2$) was calculated as the product of applied irradiance (mW/cm$^2$), duty cycle, and stimulation duration (240 s).

### S3 MR thermometry

To evaluate potential temperature changes near the site of irradiation during tPBM, MR thermometry data were collected at the highest irradiance condition (1064 nm, 200 mW/cm$^2$) for both pulsation frequencies (10 Hz and 40 Hz) across 30 subjects. As the tPBM laser was positioned over the right forehead to stimulate the right prefrontal cortex (rPFC), this region served as the target region of interest (ROI) for thermometry measurements (**Fig. S2**). Group-level statistical testing was performed using paired-sample t-tests for each pulsation frequency, comparing the mean temperature during the baseline period (Min 1-4) to the stimulation period (Min 4-8).

As shown in **Table S1**, the average temperature change was minimal and remained stable over the entire 12-minute recording. This confirmed that even at the longest wavelength and highest irradiance, no measurable heating effects were produced in the targeted brain region. As the lasers were secured such that participants were unable to see any light, and no thermal sensations were reported, both thermal and placebo effects were minimized.

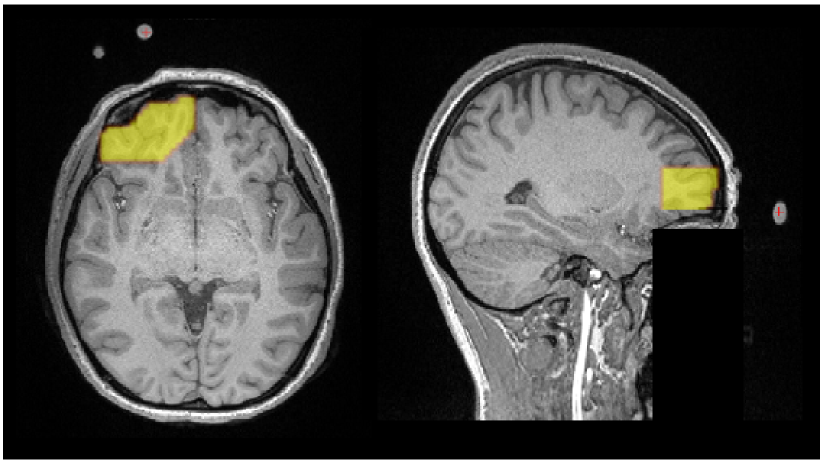

**Figure S2. The region of interest for thermometry assessment.** Vitamin E capsules were used as fiducial markers to indicate the laser location. The illuminated region encompasses approximately 3.59 cm from the site of light irradiation. This region was segmented for each participant and used for MR thermometry analysis.

**Table S1.** Group-level statistical comparison of temperature changes between the baseline and tPBM stimulation periods for both pulsation frequencies.

| Frequency | Baseline (°C) | tPBM (°C) | p |
|---|---|---|---|
| 10 Hz | -0.016 | 0.050 | 0.13 |
| 40 Hz | -0.0020 | 0.0087 | 0.76 |

#### S4 Study design and iPBM protocol

The tPBM recordings analyzed in this manuscript were collected as part of a larger study that also included intranasal photobiomodulation (iPBM) recordings. Within each session, tPBM and iPBM recordings were acquired in a fixed alternating order across eight consecutive EEG recordings. Each participant completed four tPBM and four iPBM recordings per session (**Table S2**). For the iPBM recordings, NIR light was administered through the nasal cavity via a nosepiece clipped to the right nostril, targeting the rPFC via a 10-metre, 400 um fibre cable and two Class 3 laser systems (MDL-III-808-1W and MDL-III-1064-1W; Vielight Inc., Toronto, Canada). iPBM recordings used the same two wavelengths (808 nm, 1064 nm) and two pulsation frequencies (10 Hz, 40 Hz), but at lower irradiances (5, 7, 9 mW/cm$^2$) appropriate for intranasal delivery. Each iPBM recording followed the same 12-minute PRE-DURING-POST block design (4 minutes each). The iPBM results will be reported in a separate manuscript.

This study design is relevant to the interpretation of the carry-over analysis (**Model B**; **Table S7**), which tested whether the dose level of the immediately preceding recording predicted the current tPBM response. Because tPBM and iPBM recordings were alternated, the “previous dose” for a given tPBM recording was considered as the recording that immediately preceded it, which was an iPBM recording. Similarly, the carry-over model also tested whether a preceding tPBM dose affected subsequent iPBM outcomes; these iPBM results are included in **Table S7** for completeness.

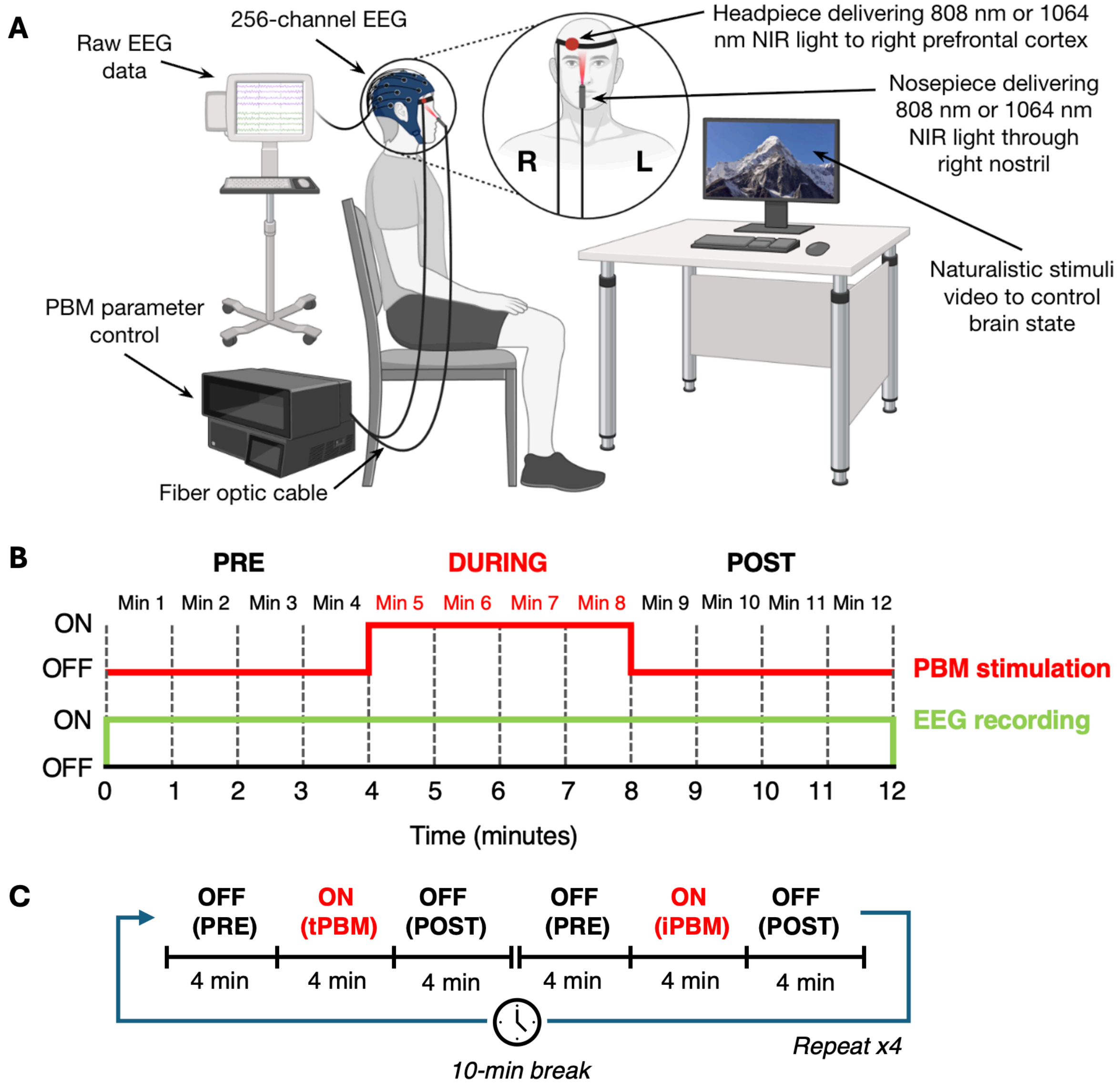


**Figure S3. Experimental setup and recording timeline.** (A) Schematic illustration of EEG-tPBM experimental setup (*BioRender.com*). (B) EEG recording timeline following block tPBM stimulus design.

**Table S2. Recording order and modality assignment within each experimental session.** Participants were randomly assigned to one of three protocols, which determined the wavelength, pulsation frequency, and irradiance used in each recording.

| Recording | Route | Protocol 1 | Protocol 2 | Protocol 3 |
|---|---|---|---|---|
| 1 | tPBM | 1064 nm, 100 mW/cm$^2$, 10 Hz | 1064 nm, 150 mW/cm$^2$, 10 Hz | 1064 nm, 150 mW/cm$^2$, 40 Hz |
| 2 | iPBM | 808 nm, 9 mW/cm$^2$, 10 Hz | 808 nm, 5 mW/cm$^2$, 10 Hz | 808 nm, 5 mW/cm$^2$, 40 Hz |
| 3 | tPBM | 808 nm, 100 mW/cm$^2$, 10 Hz | 808 nm, 150 mW/cm$^2$, 10 Hz | 808 nm, 150 mW/cm$^2$, 40 Hz |
| 4 | iPBM | 1064 nm, 9 mW/cm$^2$, 10 Hz | 1064 nm, 5 mW/cm$^2$, 10 Hz | 1064 nm, 5 mW/cm$^2$, 40 Hz |
| 5 | tPBM | 1064 nm, 200 mW/cm$^2$, 10 Hz | 1064 nm, 200 mW/cm$^2$, 40 Hz | 1064 nm, 100 mW/cm$^2$, 40 Hz |
| 6 | iPBM | 808 nm, 7 mW/cm$^2$, 10 Hz | 808 nm, 7 mW/cm$^2$, 40 Hz | 808 nm, 9 mW/cm$^2$, 40 Hz |

| | | | | |
|---|---|---|---|---|
| 7 | tPBM | 808 nm, 200 mW/cm$^2$, 10 Hz | 808 nm, 200 mW/cm$^2$, 40 Hz | 808 nm, 100 mW/cm$^2$, 40 Hz |
| 8 | iPBM | 1064 nm, 7 mW/cm$^2$, 10 Hz | 1064 nm, 7 mW/cm$^2$, 40 Hz | 1064 nm, 9 mW/cm$^2$, 40 Hz |
| **M / F** | | 8 M / 8 F | 7 M / 7 F | 9 M / 7 F |

#### S5 Model A: LME model coefficients

**Table S3. Summary of the primary LME model (Model A) for tPBM-induced EEG band power changes.** Each row represents one frequency band, time window (During-tPBM or Post-tPBM), and cluster ROI (Increase or Decrease) combination for which FWER-significant spatiotemporal clusters were identified in **Section 2.4.3.1**. All models began with the same five fixed-effect predictors (Energy, Wavelength, Frequency, Sex, and ITA) and a random intercept for subject. Backward elimination with Benjamini-Hochberg FDR correction (q=0.05) was applied iteratively (see **Table S4** for the full elimination trace). “Surviving Predictors” lists the predictors retained in the final model; a blank indicates that all predictors were eliminated. N Electrodes denotes the number of electrodes in the cluster ROI used as the dependent variable. “Estimate” reflects the mean difference in percent change from the mean of the PRE period relative to the reference level (808 nm, 10 Hz, Low energy, Male, high ITA). Asterisks denote FDR-significant contrasts (q<0.05); these correspond to the results reported in **Table 4** of the main text.

| Time Window | Band | ROI | N Electrodes | Initial Predictors | Surviving Predictors | Contrast | Estimate (%) | 95% CI | SE | FDR q |
|---|---|---|---|---|---|---|---|---|---|---|
| During-tPBM | Theta | Decrease | 119 | Energy, Wavelength, Frequency, Sex, ITA | | | | | | |
| During-tPBM | Alpha | Decrease | 172 | Energy, Wavelength, Frequency, Sex, ITA | Sex | Female vs Male | -5.102 | [-10.028, -0.176] | 2.513 | 0.0424* |
| During-tPBM | Beta | Increase | 23 | Energy, Wavelength, Frequency, Sex, ITA | | | | | | |
| During-tPBM | Beta | Decrease | 124 | Energy, Wavelength, Frequency, Sex, ITA | | | | | | |
| During-tPBM | Gamma | Increase | 42 | Energy, Wavelength, Frequency, Sex, ITA | | | | | | |
| During-tPBM | Gamma | Decrease | 68 | Energy, Wavelength, Frequency, Sex, ITA | Sex, Wavelength | 1064 vs 808 nm | 7.574 | [1.603, 13.544] | 3.046 | 0.0258* |
| During-tPBM | Gamma | Decrease | 68 | Energy, Wavelength, Frequency, Sex, ITA | Sex, Wavelength | Female vs Male | 6.739 | [0.678, 12.800] | 3.092 | 0.0293* |
| Post-tPBM | Theta | Decrease | 180 | Energy, Wavelength, Frequency, Sex, ITA | Frequency | 40 vs 10 Hz | 14.869 | [0.035, 29.704] | 7.569 | 0.0495* |
| Post-tPBM | Alpha | Decrease | 161 | Energy, Wavelength, Frequency, Sex, ITA | | | | | | |
| Post-tPBM | Beta | Decrease | 120 | Energy, Wavelength, Frequency, Sex, ITA | Sex | Female vs Male | 9.215 | [0.426, 18.003] | 4.484 | 0.0399* |
| Post-tPBM | Gamma | Increase | 45 | Energy, Wavelength, Frequency, Sex, ITA | Frequency | 40 vs 10 Hz | -36.894 | [-70.740, -3.048] | 17.269 | 0.0326* |

#### S6 Model A: Backward elimination trace

**Table S4. Backward elimination trace for the primary linear mixed-effects model (Model A, tPBM).** At each step, Benjamini-Hochberg FDR correction (q=0.05) was applied simultaneously to all fixed-effect p-values in the current model. The predictor whose contrasts yielded the highest FDR-corrected q-value was removed if that value exceeded 0.05. Elimination continued until all remaining predictors were FDR-significant or no predictors remained. Each row documents one removal step for a given frequency band, time window, and cluster ROI combination. The "FDR q-value" column reports the worst (highest) FDR-corrected q-value among the removed predictor's contrasts at that step. Predictor abbreviations: Energy = irradiance level (Low: 100, Mid: 150, High: 200 mW/cm$^2$); Wavelength = 808 or 1064 nm; Frequency = pulsation frequency (10 or 40 Hz); Sex = Male or Female; ITA = z-scored Individual Typology Angle (skin tone).

| Band | Time Window | ROI | Step | Removed Predictor | FDR q-value | Predictors Before Removal | Predictors After Removal |
|---|---|---|---|---|---|---|---|
| Theta | During-tPBM | Decrease | 1 | Energy | 0.8048 | Energy, Wavelength, Frequency, Sex, ITA | Wavelength, Frequency, Sex, ITA |
| Theta | During-tPBM | Decrease | 2 | Wavelength | 0.8015 | Wavelength, Frequency, Sex, ITA | Frequency, Sex, ITA |
| Theta | During-tPBM | Decrease | 3 | Sex | 0.8009 | Frequency, Sex, ITA | Frequency, ITA |
| Theta | During-tPBM | Decrease | 4 | ITA | 0.7717 | Frequency, ITA | Frequency |
| Theta | During-tPBM | Decrease | 5 | Frequency | 0.1274 | Frequency | |
| Alpha | During-tPBM | Decrease | 1 | Energy | 0.9857 | Energy, Wavelength, Frequency, Sex, ITA | Wavelength, Frequency, Sex, ITA |
| Alpha | During-tPBM | Decrease | 2 | Wavelength | 0.9525 | Wavelength, Frequency, Sex, ITA | Frequency, Sex, ITA |
| Alpha | During-tPBM | Decrease | 3 | Frequency | 0.8603 | Frequency, Sex, ITA | Sex, ITA |
| Alpha | During-tPBM | Decrease | 4 | ITA | 0.198 | Sex, ITA | Sex |
| Beta | During-tPBM | Increase | 1 | Energy | 0.9566 | Energy, Wavelength, Frequency, Sex, ITA | Wavelength, Frequency, Sex, ITA |
| Beta | During-tPBM | Increase | 2 | Wavelength | 0.9794 | Wavelength, Frequency, Sex, ITA | Frequency, Sex, ITA |
| Beta | During-tPBM | Increase | 3 | Frequency | 0.9795 | Frequency, Sex, ITA | Sex, ITA |
| Beta | During-tPBM | Increase | 4 | Sex | 0.9774 | Sex, ITA | ITA |
| Beta | During-tPBM | Increase | 5 | ITA | 0.9165 | ITA | |

| Band | Time Window | ROI | Step | Removed Predictor | FDR q-value | Predictors Before Removal | Predictors After Removal |
|---|---|---|---|---|---|---|---|
| Beta | During-tPBM | Decrease | 1 | ITA | 0.9979 | Energy, Wavelength, Frequency, Sex, ITA | Energy, Wavelength, Frequency, Sex |
| Beta | During-tPBM | Decrease | 2 | Energy | 0.7202 | Energy, Wavelength, Frequency, Sex | Wavelength, Frequency, Sex |
| Beta | During-tPBM | Decrease | 3 | Frequency | 0.5947 | Wavelength, Frequency, Sex | Wavelength, Sex |
| Beta | During-tPBM | Decrease | 4 | Wavelength | 0.2845 | Wavelength, Sex | Sex |
| Beta | During-tPBM | Decrease | 5 | Sex | 0.0657 | Sex | |
| Gamma | During-tPBM | Increase | 1 | Energy | 0.9287 | Energy, Wavelength, Frequency, Sex, ITA | Wavelength, Frequency, Sex, ITA |
| Gamma | During-tPBM | Increase | 2 | Frequency | 0.9281 | Wavelength, Frequency, Sex, ITA | Wavelength, Sex, ITA |
| Gamma | During-tPBM | Increase | 3 | Sex | 0.9406 | Wavelength, Sex, ITA | Wavelength, ITA |
| Gamma | During-tPBM | Increase | 4 | ITA | 0.9627 | Wavelength, ITA | Wavelength |
| Gamma | During-tPBM | Increase | 5 | Wavelength | 0.1572 | Wavelength | |
| Gamma | During-tPBM | Decrease | 1 | Frequency | 0.7362 | Energy, Wavelength, Frequency, Sex, ITA | Energy, Wavelength, Sex, ITA |
| Gamma | During-tPBM | Decrease | 2 | ITA | 0.67 | Energy, Wavelength, Sex, ITA | Energy, Wavelength, Sex |
| Gamma | During-tPBM | Decrease | 3 | Energy | 0.4825 | Energy, Wavelength, Sex | Wavelength, Sex |
| Theta | Post-tPBM | Decrease | 1 | Energy | 0.9534 | Energy, Wavelength, Frequency, Sex, ITA | Wavelength, Frequency, Sex, ITA |
| Theta | Post-tPBM | Decrease | 2 | ITA | 0.8426 | Wavelength, Frequency, Sex, ITA | Wavelength, Frequency, Sex |
| Theta | Post-tPBM | Decrease | 3 | Wavelength | 0.4891 | Wavelength, Frequency, Sex | Frequency, Sex |
| Theta | Post-tPBM | Decrease | 4 | Sex | 0.4879 | Frequency, Sex | Frequency |
| Alpha | Post-tPBM | Decrease | 1 | Energy | 0.8217 | Energy, Wavelength, Frequency, Sex, ITA | Wavelength, Frequency, Sex, ITA |
| Alpha | Post-tPBM | Decrease | 2 | Wavelength | 0.8215 | Wavelength, Frequency, Sex, ITA | Frequency, Sex, ITA |
| Alpha | Post-tPBM | Decrease | 3 | Frequency | 0.6914 | Frequency, Sex, ITA | Sex, ITA |

| Band | Time Window | ROI | Step | Removed Predictor | FDR q-value | Predictors Before Removal | Predictors After Removal |
|---|---|---|---|---|---|---|---|
| Alpha | Post-tPBM | Decrease | 4 | Sex | 0.562 | Sex, ITA | ITA |
| Alpha | Post-tPBM | Decrease | 5 | ITA | 0.6104 | ITA | |
| Beta | Post-tPBM | Decrease | 1 | Energy | 0.583 | Energy, Wavelength, Frequency, Sex, ITA | Wavelength, Frequency, Sex, ITA |
| Beta | Post-tPBM | Decrease | 2 | Frequency | 0.6091 | Wavelength, Frequency, Sex, ITA | Wavelength, Sex, ITA |
| Beta | Post-tPBM | Decrease | 3 | ITA | 0.4745 | Wavelength, Sex, ITA | Wavelength, Sex |
| Beta | Post-tPBM | Decrease | 4 | Wavelength | 0.1606 | Wavelength, Sex | Sex |
| Gamma | Post-tPBM | Increase | 1 | Energy | 0.9881 | Energy, Wavelength, Frequency, Sex, ITA | Wavelength, Frequency, Sex, ITA |
| Gamma | Post-tPBM | Increase | 2 | Sex | 0.4497 | Wavelength, Frequency, Sex, ITA | Wavelength, Frequency, ITA |
| Gamma | Post-tPBM | Increase | 3 | ITA | 0.4837 | Wavelength, Frequency, ITA | Wavelength, Frequency |
| Gamma | Post-tPBM | Increase | 4 | Wavelength | 0.1417 | Wavelength, Frequency | Frequency |

#### S7 Model B: Carry-over analysis

**Table S5. Results of the carry-over analysis (Model B).** This model tested whether the energy level of the immediately preceding recording session (PrevEnergyLevel; reference: Low) predicted the current session's EEG band power response, with subject as a random intercept. Backward elimination with FDR correction was applied identically to Model A. Because tPBM and iPBM recordings were alternated within each session (see ***Section S4***), preceding sessions include both tPBM and iPBM doses. Results are shown for both tPBM and iPBM outcomes to provide a complete picture of potential cross-modality carry-over effects.

| Modality | Time Window | Band | ROI | N Electrodes | Contrast | Estimate (%) | 95% CI | SE | FDR q |
|---|---|---|---|---|---|---|---|---|---|
| tPBM | During-tPBM | Theta | Decrease | 119 | | | | | |
| tPBM | During-tPBM | Alpha | Decrease | 172 | | | | | |
| tPBM | During-tPBM | Beta | Increase | 23 | | | | | |
| tPBM | During-tPBM | Beta | Decrease | 124 | | | | | |
| tPBM | During-tPBM | Gamma | Increase | 42 | | | | | |
| tPBM | During-tPBM | Gamma | Decrease | 68 | | | | | |
| tPBM | Post-tPBM | Theta | Decrease | 180 | | | | | |
| tPBM | Post-tPBM | Alpha | Decrease | 161 | | | | | |
| tPBM | Post-tPBM | Beta | Decrease | 120 | | | | | |
| tPBM | Post-tPBM | Gamma | Increase | 45 | | | | | |
| iPBM | During-iPBM | Theta | Increase | 6 | | | | | |
| iPBM | During-iPBM | Theta | Decrease | 79 | | | | | |
| iPBM | During-iPBM | Beta | Increase | 42 | | | | | |
| iPBM | During-iPBM | Beta | Decrease | 75 | | | | | |
| iPBM | During-iPBM | Gamma | Increase | 48 | | | | | |

| Modality | Time Window | Band | ROI | N Electrodes | Contrast | Estimate (%) | 95% CI | SE | FDR q |
|---|---|---|---|---|---|---|---|---|---|
| iPBM | During-iPBM | Gamma | Decrease | 74 | Previous High vs Low dose | -7.804 | [-14.626, -0.982] | 3.481 | 0.0250* |
| iPBM | During-iPBM | Gamma | Decrease | 74 | Previous Mid vs Low dose | -9.116 | [-15.903, -2.328] | 3.463 | 0.0170* |
| iPBM | Post-iPBM | Theta | Decrease | 156 | | | | | |
| iPBM | Post-iPBM | Beta | Increase | 32 | | | | | |
| iPBM | Post-iPBM | Beta | Decrease | 78 | | | | | |
| iPBM | Post-iPBM | Gamma | Increase | 43 | | | | | |
| iPBM | Post-iPBM | Gamma | Decrease | 66 | | | | | |

As shown in **Table S5**, For tPBM outcomes, no significant carry-over effects were detected in any band, phase, or ROI direction. For iPBM outcomes, significant carry-over effects were detected only in the gamma band during stimulation within the negative (power decrease) ROI: preceding sessions with Mid or High energy levels were associated with significantly more pronounced gamma suppression compared to Low energy.
